# The mechanism of mRNA activation

**DOI:** 10.1101/2023.11.15.567265

**Authors:** Riley C. Gentry, Nicholas A. Ide, Victoria M. Comunale, Erik W. Hartwick, Colin D. Kinz-Thompson, Ruben L. Gonzalez

## Abstract

During translation initiation, messenger RNA molecules must be identified and activated for loading into a ribosome. In this rate-limiting step, the heterotrimeric protein eukaryotic initiation factor eIF4F must recognize and productively interact with the 7-methylguanosine cap at the 5’ end of the messenger RNA and subsequently activate the message. Despite its fundamental, regulatory role in gene expression, the molecular events underlying cap recognition and messenger RNA activation remain mysterious. Here, we generate a unique, single-molecule fluorescence imaging system to interrogate the dynamics with which eIF4F discriminates productive and non-productive locations on full-length, native messenger RNA molecules. At the single-molecule level, we observe stochastic sampling of eIF4F along the length of the messenger RNA and identify allosteric communication between the eIF4F subunits which ultimately drive cap-recognition and subsequent activation of the message. Our experiments uncover novel functions for each subunit of eIF4F and we conclude by presenting a model for messenger RNA activation which precisely defines the composition of the activated message. This model provides a general framework for understanding how messenger RNA molecules may be discriminated from one another, and how other RNA-binding proteins may control the efficiency of translation initiation.

## Introduction

The different efficiencies with which the ribosome translates individual messenger RNAs (mRNAs) is a tightly regulated, critical component of gene expression. In eukaryotes, much of this regulation is dictated during mRNA activation, whereby mRNA molecules are primed for loading into a ribosomal 43S pre-initiation complex (PIC)^1–3^. More than 95% of cellular mRNAs are activated through a 5’ 7-methylguanosine cap (cap)-dependent process^4,5^ driven by the highly conserved eukaryotic initiation factor (eIF) 4F, a heterotrimeric complex composed of eIF4A, eIF4E and eIF4G^1–3^. Due to the low abundance of eIF4G and regulated sequestration of eIF4E, mRNA activation is the rate-limiting step of translation for most genes^1,6^. Consequently, healthy cells actively adjust the pool of free eIF4E to drive normal changes in gene expression, while many viruses and diseased cells rely on either physically altered^7^ or stoichiometrically dysregulated^8^ eIF4F to proliferate.

Despite its prominent role in cellular homeostasis, and decades of biochemical research^4–6,9–32^, the molecular mechanism by which eIF4F activates mRNAs remains completely mysterious. Prior studies have assigned roles for each component of eIF4F: eIF4E is a small cap-binding protein; eIF4A is a DEAD-Box RNA helicase; and eIF4G serves as the core component of eIF4F and a scaffolding protein from which eIF4E and eIF4A exert their activity, while also likely bridging the mRNA to the PIC^1–3^. Using this information, these studies have culminated in a working model for mRNA activation^1–3,33^ by which eIF4E serves as the initial point of contact between eIF4F and the mRNA, near the cap in the 5’ untranslated region (UTR) of the mRNA. This contact is then stabilized by the RNA binding activity of eIF4G. Finally, eIF4A hydrolyzes ATP to unwind RNA secondary structure elements near the cap and results in additional, unknown steps which transition the mRNA from an ‘inactivated’ state to an ‘activated’ state.

Notably, there exists a plethora of evidence contradicting the model described above^2,3,5,11,15,16,19,21,25,28,30,32–34^, and neither the molecular basis for the preferential association of eIF4F with the 5’ UTR of the mRNA, nor the identity of the ‘activated’ state of the mRNA have been experimentally determined. These questions remain open, in part, due to the technical limitations of working with eIF4G. Recombinant full-length eIF4G is notoriously difficult to purify and possesses potent non-specific RNA binding activity^30,35^. Consequently, many biochemical studies have been performed with truncated eIF4G constructs^14,15,19,23,24,26,36–39^ together with short, minimal-length mRNA mimics to minimize nonspecific binding^15,24,30,38,39^. Biochemical studies have also been hampered by the lack of a complete structure of eIF4F^13,14,27,38^. Moreover, no direct assay for mRNA activation has been developed and instead assays which measure ribosome stalling on the start codon, downstream of mRNA activation and subsequent 43S PIC loading, are used to indirectly infer eIF4F function^30,35^.

To overcome these limitations and determine the mechanism of mRNA activation, here we have developed a single-molecule fluorescence resonance energy transfer (smFRET) system which probes the binding of yeast eIF4F to local segments of a native, full-length mRNA. To achieve this, we site-specifically labeled native mRNAs at either cap-proximal or cap-distal locations, and site-specifically labeled full-length eIF4G, allowing us to systematically interrogate the binding kinetics of eIF4F and eIF4F-subcomplexes at cap-proximal and cap-distal locations on full-length mRNA. By comparing the rates determined by our experiments, which span >30,000 individual molecules queried across >60 experimental conditions, we uncover the molecular basis for cap recognition and ascribe new functions to each subunit of eIF4F. By directly following eIF4G, we isolate an intermediate in mRNA activation and define the ‘activated’ state of the mRNA. These insights allow us to conclude by presenting a model for mRNA activation, which implies new ways cells likely regulate gene expression.

### Direct observation of eIF4G binding dynamics

To develop a system allowing us to monitor mRNA binding by eIF4F and eIF4F subcomplexes, we first reconstituted *Saccharomyces cerevisiae* translation initiation following established protocols (**Extended Data Figure 1a**)^30,35,40,41^. Notably, we optimized the purification procedure for eIF4G, enabling us to purify large quantities of full-length eIF4G virtually free of degradation products, which are otherwise common contaminants in eIF4G preparations^30,35^. We validated the function of recombinant eIF4F using a standard native gel-shift assays with a capped *S. cerevisiae* mRNA encoding for the gene rpl41a (henceforth, rpl41a)^30,35^. This assay measures the ability of 43S PICs to load onto and scan to the start codon of capped mRNAs. As expected, efficient ribosome recruitment was dependent on eIF4F, with reactions lacking eIF4E but containing eIF4G and eIF4A showing diminished but detectable recruitment (**Extended Data Figure 1b**)^30^. Next, we performed titration-regime ensemble binding measurements using fluorescence anisotropy to measure the binding behavior of eIF4G with a short, uncapped 51-nucleotide fragment of rpl41a containing the 5’ untranslated region (UTR) and first 9 codons^42^. Consistent with previous publications, we observed that eIF4G associated tightly^30^ and multivalently^5^ with the short, uncapped mRNA fragment (**Extended Data Figure 1c**).

To assess the validity of a model whereby eIF4E serves as the first point of contact between eIF4F and the mRNA 5’ UTR, one must compare the binding behavior of eIF4F at cap proximal and cap distal locations in the absence and presence of the cap and eIF4E. The robust, nonspecific RNA binding activity of eIF4G complicates measurement of eIF4F association with the cap because most assays are agnostic to binding location. We therefore sought to develop an assay sensitive to the binding location of eIF4F on full-length mRNAs using a series of smFRET constructs to probe the association and dissociation kinetics of both eIF4F and eIF4F-subcomplexes to cap-proximal and cap-distal locations on full-length rpl41a. We site-specifically labeled rpl41a using a deoxyribozyme^43^ and selectively capped the labeled mRNAs to generate three constructs: (1) capped rpl41a with a cap-proximal Cy3 FRET donor fluorophore located at nucleotide 14 on the mRNA (capped), (2) uncapped rpl41a with a cap-proximal Cy3 (uncapped), and (3) capped rpl41a with a cap-distal Cy3 located at position 92 on the mRNA (internal) (**Figure 1a**). Next, we site-specifically labeled eIF4G with the Cy5 FRET acceptor fluorophore the linker region between the eIF4E binding motif and the second RNA binding motif (**Extended Data Figure 1a**). The Cy3- and Cy5-labeled mRNA and protein pairs allow us to directly observe the binding kinetics of any Cy5-eIF4G-containing complex to localized cap-proximal or cap-distal positions on full-length Cy3-rpl41a mRNA (Cy3-rpl41a).

**Figure 1.**
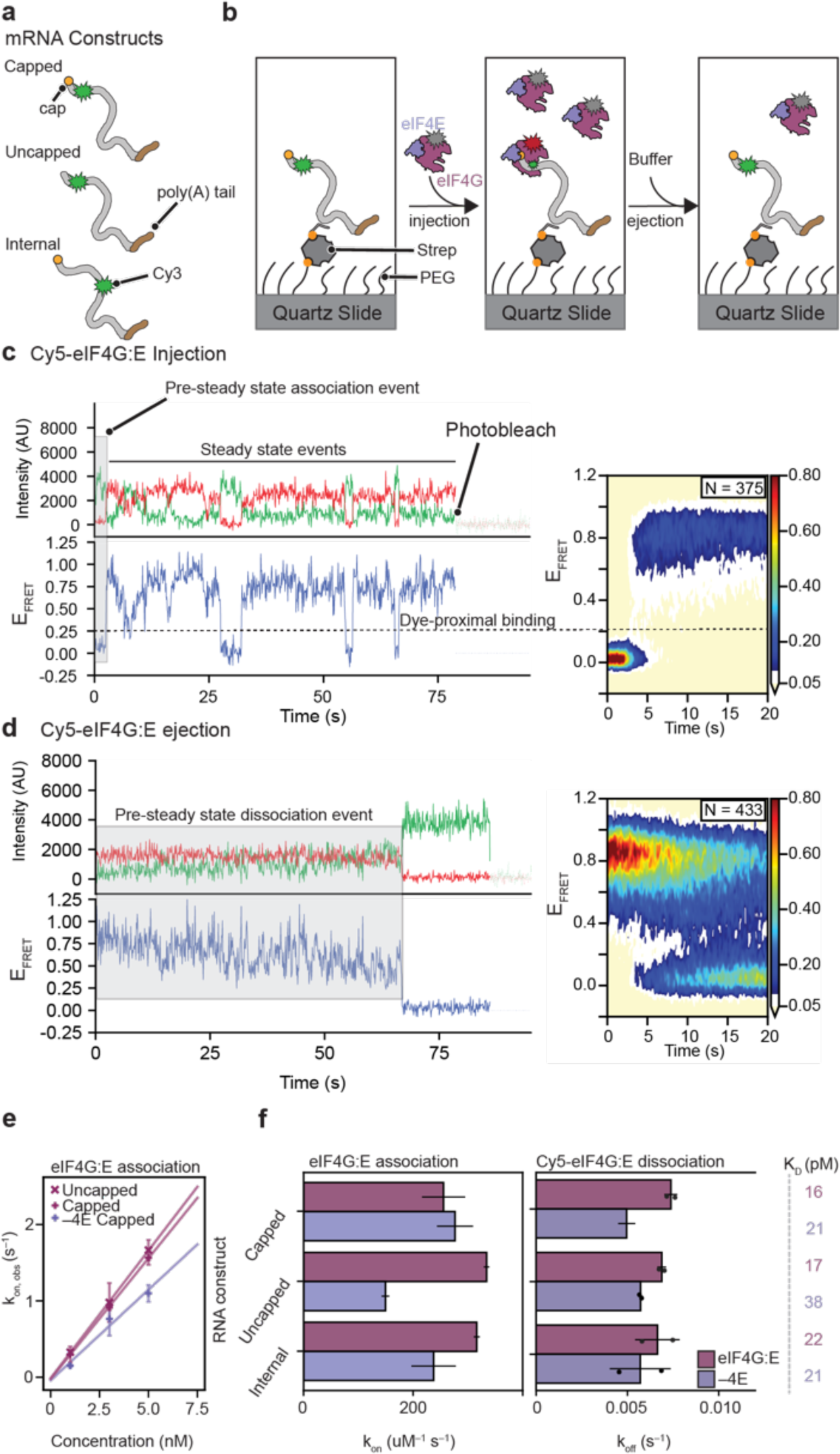
eIF4G/G:E stably associates with mRNA independent of its binding position or the cap. (a) Cartoon schematic of the full-length, Cy3-labeled rpl41a mRNA constructs used in this work. (b) Cartoon representation of the stopped flow injection and ejection pre-steady state experiments used to measure kon and koff. (c) An example trace taken from a Cy5-eIF4G:E injection experiment is displayed. The trace can be divided into two portions: the first binding event in the trajectory encompasses the pre-steady-state portion, whereas the later portions of the trajectory display steady-state binding and dissociation events until bleaching of the donor fluorophore. On the right, a surface contour plot showing the aggregate behavior of all the trajectories in an injection experiment is displayed. (d) The equivalent example trace and surface contour plot for ejection experiments are shown. Since all of the excess Cy5-eIF4G:E has been removed from the flowcell, only the pre-steady-state behavior can be inferred from ejection experiments. (e) The association rate of Cy5-eIF4G/G:E is linear with respect to concentration and depends more on the presence of eIF4E than on the cap. (f) Quantitation of the pre-steady state association and dissociation rates for Cy5-eIF4G/G:E, demonstrating that eIF4G uniformly binds to mRNA irrespective of the cap or its binding location.

We performed our experiments using a wide-field, prism-based, total internal reflection (TIR) fluorescence microscope, which enables the collection of movies containing information from hundreds of individual molecules simultaneously (**Supplemental Movie 1**). To perform these single-molecule experiments, we sparsely tether individual Cy3-rpl41a mRNAs hybridized at an internal position in the mRNA with a short biotinylated DNA-oligonucleotide to the surface of a quartz flowcell passivated with a mixture of methoxy-terminated polyethylene glycol (PEG) and biotin-terminated PEG via a biotin-streptavidin bridge (**Figure 1b**). We can inject and freely exchange solution in the flowcell before and during experiments, enabling both steady-state and pre-steady-state measurements. After tethering the Cy3-rpl41a mRNA, we inject Cy5-eIF4G containing complexes into the flowcell, and directly excite Cy3 with TIR laser light. When Cy5-eIF4G binds the mRNA at a position within ∼7 nm of the Cy3 on the Cy3-rpl41a, FRET causes the Cy3 emission to decrease while stimulating Cy5 emission (**Extended Data Figure 2a,b**). Although the real power of single-molecule experiments comes from the analysis of individual trajectories, many of the conclusions in this work can be surmised upon viewing the unprocessed movies (**Supplemental Movies S1-S7**).

Before performing experiments on full-length rpl41a mRNA, we first validated our choice of labeling constructs by performing smFRET experiments between Cy5-eIF4G co-purified as a complex with eIF4E (Cy5-eIF4G:E), and the capped, 24-nucleotide 5’ UTR of rpl41a with the cap-proximal Cy3 labeling position, which we expected to only bind a single eIF4G molecule (**Extended Data Figure 2a,b**). These experiments confirm robust FRET between the Cy3-mRNA fragment and Cy5-eIF4G:E and reveal a rapid, but transient association of Cy5-eIF4G:E with the 5’ UTR (**Supplemental Movie S1, Extended Data Figure 2c**). Unexpectedly, both the association and dissociation rates of eIF4G:E scaled with increasing concentration (**Extended Data Figure 2d**). Concentration-dependent dissociation rates are indicative of facilitated dissociation, a phenomenon that has been observed for multiple nucleic-acid binding proteins that make multivalent interactions with their substrates^44–48^. Because eIF4G possesses three RNA binding motifs, at any given time one or more of these motifs may dissociate from the mRNA while the others remain bound, thus preventing eIF4G from being released from the mRNA. However, when an individual binding motif releases from the mRNA, the vacancy can be filled by another competing eIF4G molecule which in turn facilitates the dissociation of the original eIF4G molecule (**Extended Data Figure 2e**).

### eIF4G associates tightly and nonspecifically along mRNA

Having validated the smFRET constructs, we sought to understand the molecular basis for 5’ end recognition by eIF4F. Since eIF4G provides most of eIF4F’s binding stability to mRNA and eIF4E directly associates with the cap^5,16,20,25,30^, we hypothesized that we could determine the basis for the assumed preferential association of eIF4F with the mRNA 5’ end by measuring the association and dissociation rates of Cy5-eIF4G and Cy5-eIF4G:E to the capped, uncapped, and internal constructs (**Figure 1a**) (cases where identical experiments containing either Cy5-eIF4G or Cy5-eIF4G:E were performed will henceforth be denoted Cy5-eIF4G/G:E). To determine association (k_on_) and dissociation (k_off_) rate constants, we performed pre-steady-state measurements. To measure association rates, we delivered Cy5-eIF4G/G:E to surface tethered Cy3-rpl41a while initiating data collection (**Figure 1b, injection, Supplemental Movies 2,3**). The waiting time between the delivery of Cy5-eIF4G or Cy5-eIF4G:E and the first binding event is indicative of the association rate in pre-steady-state conditions, while the apparent association and dissociation rates in steady-state can be inferred from the later timepoints for each individual molecule (**Figure 1c**). To measure dissociation rates, we started with Cy3-mRNA molecules pre-bound to Cy5-eIF4G/G:E, and then removed all free protein by flushing the flowcell with buffer lacking initiation factors (**Figure 1b, ejection**), and measured the time required for Cy5-eIF4G/G:E dissociation (**Figure 1d**). Cy5-eIF4G/G:E binding to the mRNA resulted in a broad distribution of FRET efficiencies (E_FRET_), likely due to eIF4G binding at slightly different positions along the mRNA (**Figure 1c,d right**). Therefore, to prevent overinterpretation of the different E_FRET_ values, we simplified our analysis to only classify E_FRET_ states as either “bound” or “unbound”, corresponding to high and low (near zero) E_FRET_ values (**Figure 1c,d**).

We measured the association rates between Cy5-eIF4G/G:E and each mRNA construct. If cap-recognition by eIF4E serves as the initial point of contact between eIF4F and the mRNA, one would predict Cy5-eIF4G:E to most rapidly associate to cap-proximal positions, while association of Cy5-eIF4G in the absence of eIF4E would not be expected to display any cap dependence. To our surprise, eIF4G associated rapidly with the mRNA in all experiments, with association rate constants ranging from 150-300 µM^−1^ s^−1^, which was independent of the cap, and the binding position of eIF4G (**Figure 1e**). We observed a slight decrease in the association of Cy5-eIF4G to cap-proximal positions in the absence of eIF4E, but this effect was not cap-dependent (**Figure 1e**). These measurements show that any additional interaction between eIF4E and the cap does not allow eIF4G:E to more rapidly associate with cap-proximal positions on mRNA.

Having observed little variation in the association rates of eIF4G, we expected eIF4E to selectively stabilize eIF4G:E at the cap. Surprisingly, eIF4G dissociated very slowly from all three mRNA constructs both with and without eIF4E (**Figure 1f, Extended Data Fig 3**). Our initial experiments measured dissociation rates slow enough that they were limited by photobleaching, so we performed experiments where we shuttered the excitation laser every 3 seconds to better preserve the fluorophores, and, under these conditions, Cy5-eIF4G and Cy5-eIF4G:E remained bound for upwards of minutes (**Figure 1f**). Notably, we also performed an experiment with 30 second shuttering intervals which showed stable eIF4G:E association for tens of minutes (**Supplemental Movie 4**), thus the k_off_ values we observed (**Figure 1f**) likely represent upper-bounds for the true dissociation rate of eIF4G from the mRNA. These dissociation rates may also not necessarily reflect true dissociation rates from the mRNA molecule because facilitated dissociation may allow for intra-mRNA “hopping”^46^, which we also observe (**Extended Data Figure 4**). Although upper-bounds, these k_off_ values are still informative because they provide upper limits for the equilibrium dissociation constants (K_D_s) of eIF4G/G:E in the range of 10-50 pM for each mRNA construct **(Figure 1f)**.

Variations in the affinity of eIF4G/G:E across productive, capped, locations on the mRNA and non-productive, uncapped and internal positions, are minor. These minor differences cannot be reconciled with a model for mRNA activation that begins with preferential association of eIF4F at the 5’ cap. The lack of cap-dependence suggests the functional consequence of cap recognition may occur at a step downstream of initial binding by eIF4G:E^5^. However, the cellular concentration of eIF4G is roughly stoichiometric to that of mRNA molecules^6^, and each molecule contains multiple potential non-specific binding sites. Since eIF4G dissociates so slowly from these cap-distal, non-productive locations on the mRNA, it is unlikely that eIF4G:E simply passively samples the mRNA and only selectively functions when bound to cap. Although eIF4G can be stably co-purified with eIF4E^18,30,35^, another possibility is that eIF4E dissociates from RNA-bound eIF4G unless the cap is nearby to provide additional stability, thus the absence or presence of eIF4E may distinguish productive and nonproductive eIF4G.

### eIF4G:E associates with mRNA as a complex lacking cap specificity

Because we placed the acceptor fluorophore on eIF4G, we were not able to distinguish if eIF4E was present when eIF4G bound to the mRNA. To ascertain whether eIF4E was present when eIF4G:E was bound to each of the capped, uncapped, and internal mRNA constructs, we utilized a site-specifically Cy5-labeled eIF4E (Cy5-eIF4E) that has been characterized^16,20,25^. Consistent with the literature^16,20,25^, Cy5-eIF4E showed very transient sampling of the cap (**Figure 2a, Supplemental Movie 5, left**), and did not show detectable binding to the uncapped or internal constructs. Similar to Cy5-eIF4G:E, when we utilized eIF4G:E-Cy5, generated by supplying unlabeled eIF4G to Cy5-eIF4E, the steady-state time between association events was reduced and the duration of binding events was lengthened by nearly an order of magnitude (**Figure 2b, Supplemental Movie 5, right**)^16,20,25^, but the complex still facilitated its own dissociation (**Extended Data Figure 5**). Unexpectedly, but in agreement with our Cy5-eIF4G:E experiments, eIF4G:E-Cy5 showed no cap- or positional-dependence on association or dissociation in either steady-state (**Figure 2b**) or pre-steady state measurements (**Figure 2c, Supplemental Movie 6**). The eIF4G:E association and dissociation rates for experiments using Cy5-eIF4E matched those measured in experiments using Cy5-eIF4G (**Figure 2c**), demonstrating that eIF4G and eIF4E function in concert as a complex, even when bound to cap-distal locations on mRNA. The lack of cap-specify suggests that rather than serving as the initial point of contact between eIF4F and the mRNA, cap sensing by eIF4E is likely functionally relevant downstream of the initial binding event. However, if the cap does not serve as a scaffold to assemble eIF4F at the correct location on the mRNA, as previously thought^1–3^, some other factor must direct eIF4F binding to the 5’ end.

### eIF4A drives cap-recognition

To interrogate the role of the 5’ cap during mRNA activation, we focused on eIF4A and eIF4B, which are thought to participate in mRNA activation downstream of eIF4F binding^1–3^. Utilizing Cy5-eIF4G, we assessed the binding behavior of Cy5-eIF4G:E to each of the mRNA constructs in the presence of eIF4A, eIF4B, and ATP•Mg^2+^. We began with tethered Cy3-mRNA constructs and delivered small amounts of Cy5-eIF4G:E in the presence of saturating amounts of eIF4A and ATP•Mg^2+^ (Cy5-eIF4G:E+A) or eIF4A, eIF4B and ATP•Mg^2+^ (Cy5-eIF4G:E+A+B) and measured the waiting time for the first binding event. Similar to Cy5-eIF4G:E, Cy5-eIF4G:E:A:B showed neither a cap-nor positional-dependence on its association with mRNA (**Extended Data Figure 6a**). However, the steady-state experiments in the presence of eIF4A and eIF4A+B showed cap-dependent changes in the dissociation behavior (**Extended Data Figure 6b**).

To better assess the effects of eIF4A, eIF4B, and ATP•Mg^2+^ on the dissociation rate of Cy5-eIF4G:E, we performed pre-steady state experiments. In these experiments we began with Cy5-eIF4G:E bound to the mRNA, and exchanged the buffer with new buffer containing saturating amounts of eIF4A, eIF4B, and ATP•Mg^2+^, but lacking eIF4G and eIF4E (**Figure 3, right**). We compared these experiments to the dissociation experiments performed earlier (**Figure 3, left**), in which we delivered buffer containing no additional proteins. We performed two sets of experiments: one in which the laser continuously illuminated the sample, and one in which the laser was shuttered every 3 seconds, allowing us to determine rates on both the seconds and minutes timescales (**Figure 3a-d**).

We observed that Cy5-eIF4G/G:E remains stably associated for minutes with all mRNA constructs in the absence of eIF4A, eIF4B and ATP (**Figures 3a-e**). Contrary to all expectations, in the presence of eIF4A, eIF4B, and ATP, however, binding of Cy5-eIF4G/G:E to the mRNA was destabilized by ∼30-40 fold under all conditions tested in which cap recognition was not possible (**Figure 3b-e**). In striking contrast, under the single condition where cap recognition was possible, Cy5-eIF4G:E was ∼10-fold stabilized relative to the other conditions. This order-of-magnitude increase in stability demonstrates that eIF4A discriminates the productive association of Cy5-eIF4G:E at the cap from non-productive association at cap-distal positions (**Figure 3e, Supplemental Movie 7)**. From these experiments, we conclude that eIF4A is a ‘strippase’ that removes eIF4G/G:E from non-productive locations. Moreover, we observe that cap-recognition by eIF4E inhibits this strippase activity, likely through allosteric changes in eIF4G, since eIF4E and eIF4A do not directly interact. More broadly, our experiments illustrate that cap-recognition is a multi-step process during which eIF4F must stochastically sample many binding locations and be recycled by eIF4A before arriving at the cap. Thus, the transient, but relatively long-lived eIF4F binding event at the cap is the ‘activated state’ of the mRNA. This cap-dependent stalling of eIF4F then likely serves as a beacon that facilitates the downstream loading of the PIC onto the 5’ end of the mRNA through a bridge between eIF4G and eIF3 or eIF5^10^. This conclusion is further supported by recent single-molecule observations demonstrating that pre-incubating mRNA with eIF4F before 43S PICs removes the slow step of mRNA loading in a manner dependent on the concentration of eIF4A^49^.

To test whether eIF4B or ATP were critical for eIF4G:E destabilization, we performed similar experiments in the absence of eIF4B, or in the absence of ATP (**Extended Figure 7**). In the absence of eIF4B, eIF4A was able to strip eIF4G:E from the mRNA with comparable rates as in the presence of eIF4B. However, ATP was essential for the stripping activity of eIF4A. These results reveal that eIF4A functions to remove eIF4G:E from non-productive binding locations and selectively stabilize it at the cap, thereby facilitating the search for the cap. We were curious if eIF4A might function by stepwise stripping of eIF4E from eIF4G and then eIF4G from the mRNA, so we performed similar experiments with eIF4G:E-Cy5 (**Extended figure 8**), but the dissociation rates were within two-fold of those with Cy5-eIF4G:E (**Extended Data Table 1**), suggesting that eIF4G:E is ejected as a complex.

## Discussion

How eIF4F activates mRNA for translation has remained mysterious since its discovery. Current models for mRNA activation begin with the preferential association of eIF4F with the cap-proximal 5’ UTR of the mRNA, guided by an initial contact of eIF4E with the cap^1–3,16,25^. Afterwards, eIF4F hydrolyzes ATP to exert an unknown activity which transforms the mRNA into a compositionally-undefined ‘activated’ state through a series of unknown steps^1–3^, after which some studies have suggested eIF4A recycles the complex^4,16,29,34^.

Contrary to expectations, our single-molecule experiments (summarized in **Extended Data Table 2**), define a molecular mechanism whereby eIF4E does not initially bind to the cap, but instead acts together with eIF4G and eIF4A to find the 5’ cap, thus forming the ‘activated’ state. Surprisingly, we found that eIF4G:E tightly and nonspecifically binds to locations throughout an mRNA regardless of the cap, and rather than functioning solely as a helicase, eIF4A instead strips eIF4G:E from uncapped and cap-distal positions. Although unexpected, this result is consistent with the requirement for eIF4A regardless of whether or not there is the cap-proximal secondary structure in the mRNA^11,22^ and is also consistent with some early speculations about the enzyme^33^. Perhaps even more unexpected, we found that rather than serving as the initial point of binding for eIF4F complex assembly, eIF4E recognition of the cap occurred downstream of the initial eIF4G:E binding event^5^. Cap-recognition by eIF4E allosterically inhibited eIF4A through eIF4G, preventing it from stripping eIF4G:E, thus forming the ‘activated’ state of the mRNA. Therefore because eIF4A is a notoriously poor RNA helicas that is required regardless of the mRNA secondary-structure and because eIF4G associates with other, more potent RNA helicases^50–52^, we speculate that the primary function of eIF4A during mRNA activation is that of an ATP-dependent “strippase” rather than as a helicase. Notably, other non-helicase functions have been identified for eIF4A in the human translation system^17,21^.

Collectively, our results allow us to propose a new model for mRNA activation (**Figure 4**). In the first step, eIF4G:E stochastically binds to the mRNA. In the next step, eIF4A attempts to strip eIF4G:E from the mRNA, bifurcating the cycle. If eIF4G:E is bound in a cap-distal, non-productive location, eIF4A is able to eject eIF4G:E from its current binding location by either facilitating hopping to a new location on the mRNA or releasing eIF4G:E from the mRNA entirely. However, if eIF4G:E binds near the cap and eIF4E is able to bind the cap, an eIF4E-dependent conformational change of eIF4G likely renders eIF4A:ATP unable to strip eIF4G:E, trapping the eIF4G:E:A complex at the 5’ cap. This 5’-cap-stalled eIF4F:ATP constitutes the ‘activated’ state of the mRNA. Finally, we speculate that 43S PICs can contact eIF4G:E:A on the activated mRNA and reactivate the strippase modality of eIF4A, since PICs are known to stimulate ATP hydrolysis by eIF4A^11^. This reactivation then likely facilitaties the exchange of eIF4F for the 43S PIC, which then loads onto the mRNA. Because the intracellular concentration of eIF4A is incredibly high^6^, even its modest affinity to eIF4G enables it to nearly always be bound to eIF4G:E, although it may be constantly exchanging with free eIF4A. Notably, the stochastic sampling of the mRNA by eIF4F that we observe explains early reports that 4-5 rounds of eIF4F activity were required to facilitate recruitment^3,53^ and recent *in vivo* reports that translation levels occur in oscillating bursts which can be modulated by eIF4F^54^.

While much effort has gone into looking for mRNA sequences, or proteins that regulate eIF4F binding at the 5’ end^9,15,55,56^, our results suggest cells may regulate the translation efficiency by encoding factors that accelerate the search for the cap, rather than the stability of the resulting ‘activated’ mRNA. It is also possible that cap-proximal RNA sequences, which are known to alter the affinity of eIF4E for the cap in the absence of eIF4G^57^, also alter the lifetime of the ‘activated’ state. Moreover, although our work here focused on cap-dependent translation, cap-independent translation is often eIF4G- and eIF4A dependent^7,31^. This suggests that the allosteric communication transduced by cap-bound eIF4E through eIF4G^19,32^ can be induced by other factors during non-canonical translation initiation.

## Materials and Methods

### Expression and purification of non-eIF4G protein factors

Yeast ribosomes and the yeast initiation factors eIF1, eIF1A, eIF4A, eIF4B, and eIF4E were expressed and purified from *Escherichia coli* BL21(DE3) RIPL (Agilent) following established procedures^30,35,40,41^. eIF’s 1, 1A, 4A, and 4E were expressed with N-terminal hexahistidine (6×-his) tags followed by a Tobacco Etch Virus (TEV) protease cleavage site. eIF4B was expressed with a C-terminal TEV site followed by a 6×-his tag. Native, 6×-his tagged eIF2 was expressed and purified from *S. cerevisiae* strain GP3511 as previously described^40^. Native, 6×-his-tagged eIF3 was purified from *S. cerevisiae* strain LPY87 following standard procedures with minor deviations^35^. After affinity purification, rather than phosphocellulose chromatography, eIF3 was applied to a 5mL HiTrap Q HP column (Cytiva) and its final gel filtration and storage buffer consisted of 20mM 4-(2-hydroxyethyl)-1-piperazineethanesulfonic acid (HEPES) pH=7.4, 200 mM potassium chloride (KCl), 10% glycerol, 2 mM dithiothreitol (DTT).

### Expression of eIF4G

eIF4G was expressed from a pET28a vector containing a C-terminal TEV protease cleavage site followed by an FH8 tag and 6×-his tag. This results in an ENLYQ scar on the C-terminus of eIF4G at the end of purification. In order to express eIF4G, BL21(DE3) RIPL cells were freshly transformed with pET28a-eIF4G and plated on Luria broth (LB) agar plates containing 100ug/mL carbenicillin (carb). After transformation, plates were incubated at 37 °C overnight and then kept at room temperature for up to 10 hours until inoculation of overnight cultures in 2.5 L ultra-yield flasks (Thomson Instrument Company) containing 500 mL of LB supplemented with carb, 34 µg/mL chloramphenicol, and 0.6% glucose. Overnight cultures were grown for 14-16 hours at 32 °C and then diluted 62.5:1 into 12 L of fresh LB carb + 1% glucose split across 8 ultra-yield flasks. The diluted cell culture was grown in a refrigerated shaker with a shaking speed of 215 rpm at 30°C until it reached an optical density at 600 nm (OD_600_) of ∼0.3 (approximately 2.5 hours). The shaker was then shifted to 25 °C while shaking for 1 hour. Next, eIF4G expression was induced by addition of 670 uL of freshly prepared 1 M isopropyl ß-D-1-thiogalactopyranoside (IPTG) to each flask resulting in a final IPTG concentration of ∼0.45 mM. Upon induction, the temperature of the shaker was set to 22°C and the cells were allowed to induce for 1 hour and 15 minutes. After induction, cell flasks were immediately moved to a plastic autoclave bin filled with ice and pelleted by repeated centrifugation at 5500 relative centrifugal force (rcf) for 5 minutes. Pelleted cells were resuspended in ∼35 mL of eIF4G Lysis Buffer (50 mM HEPES pH=7.4, 500 mM KCl, 10% glycerol) crudely measured in a 50mL conical tube and frozen as droplets by slowly dispensing the resuspended cell culture into liquid nitrogen. The resulting cell ‘popcorn’ can be stored in the −80 °C for at least 2 months. The cell popcorn was then lysed in a freezer mill (SPEX) using 7 cycles of 2 minutes on-time at a rate of 10 cycles per second, followed by 1 minute of off-time. The resulting cell powder was then stored at –80 °C and usable for at least 2 months.

### Purification of eIF4G

Cell lysate powder from 12 L of cell culture was resuspended in 300 mL of room temperature eIF4G Lysis Buffer supplemented with 2 EDTA-free protease inhibitor tablets (Roche), 20mM imidazole, 10mM magnesium chloride (MgCl_2_) and 50 µg of nuclease A from *Serratia Marcescens* by slowly adding the powder to actively stirring lysis buffer in a 1 L glass beaker. It takes approximately 20 minutes to resuspend the cell powder. The lysate was then clarified at 31,500 rcf for 35 minutes and the supernatant was filtered through a 0.45-micron bottle-top filter before loading onto a chelating column (Cytiva) charged with nickel and pre-equilibrated with eIF4G Lysis Buffer. eIF4G was eluted from the chelating column using a 10 column-volume (c.v.) gradient of eIF4G Elution Buffer (3:1 eIF4G Lysis Buffer: 2 M imidazole pH=8). The entire peak from the nickel column was pooled and diluted 8-fold in 20 mM HEPES-KOH pH=7 and loaded onto a HiRes Capto Q 5/50 column (Cytiva) pre-equilibrated in Low Ion-Exchange Buffer (20 mM HEPES-KOH pH=7, 100 mM KOAc, 5% glycerol) and eluted using a 20 c.v. gradient into High Ion-Exchange Buffer (20 mM HEPES-KOH pH=7, 1 M KOAc, 5% glycerol). The elution profile from the HiRes CaptoQ is very broad but monitoring absorbance at both 260 nm and 280 nm allows the fractions containing protein to be separated from contaminating RNA. The RNA-depleted, protein-containing fractions were then pooled and diluted 5-fold in 20 mM HEPES-KOH before loading onto a HiRes CaptoS 5/50 column (Cytiva) pre-equilibrated in Low Ion-Exchange Buffer and eluted across a biphasic gradient of 0-40% for 15 c.v followed by 40-100% for 7 c.v. in High Ion-Exchange Buffer. At this step, we routinely collect ∼3-4 mL of protein which elutes as a sharp peak around 31 mS/cm conductivity.

Following purification using the HiRes CaptoS, 1/100 w/w TEV protease and 4 mM beta-mercaptoethanol (BME) was added to the sample, and the sample was moved to a 4°C refrigerator. TEV protease was then allowed to cleave overnight without agitation. The following morning, 20 mM imidazole is added to the cleavage reaction, and 400uL of Ni-NTA slurry (Goldbio) equilibrated in Low Ion-Exchange Buffer was added to the sample in a 15 mL conical tube. The Ni-NTA beads and protein solution were then mixed by pipetting up and down 3 times every 3 minutes for 30 minutes at room temperature. After allowing uncleaved protein and TEV protease to bind the Ni-NTA beads, the beads and uncleaved protein are removed from the sample by passing over a gravity column. The flowthrough from the gravity column was then diluted to 35mL using Low Ion-Exchange buffer and filtered through a 0.22-micron syringe filter and loaded back onto the HiRes CaptoS column using the same conditions as earlier in the purification. A ∼1 mL elution from this gradient is then pooled and protein concentration was measured using a Bradford assay with bovine serum albumin as a standard before aliquoting into 10 µL single-use aliquots and snap freezing in liquid nitrogen.

eIF4G:E was expressed and purified identically to eIF4G with the exception that cells are transformed with both pET28a-eIF4G and a pColA-backbone containing Kanamycin resistance and full-length, untagged eIF4E under the control of a T7 promoter. Kanamycin is therefore supplemented in all of the carb-containing media described for eIF4G above.

### Fluorophore labeling

Cy5-eIF4E was prepared by purifying eIF4E-A124C and labeling with sulfo-Cy5 maleimide (Lumiprobe), as previously described^20,25^. The labeling efficiency for eIF4E was approximately 50% based on the absorbance of protein at 280 nm compared to the absorbance of the dye at 650 nm. eIF4G:E-Cy5 was generated by mixing equimolar ratios of eIF4G with Cy5-eIF4E prior to aliquoting and snap freezing in liquid nitrogen.

Cy5-eIF4G/G:E were prepared by purifying eIF4G/G:E-S501C according to the procedure above and subsequently labeling the purified protein by incubating it with a 10-fold molar excess of sulfo-Cy5 maleimide at 4C overnight. Next, the labeling mixtures were spun threw 0.22 micron benchtop spin filters (Ultrafree-MC, Millipore) and loaded onto a Superdex 200 increase 10/300 GL column (Cytiva) equilibrated in Low Ion-Exchange Buffer. The peak containing labeled eIF4G/G:E was collected and subsequently loaded onto a Capto HiResS 5/50 and further purified using the same procedure as above. eIF4G and eIF4G:E are estimated to be labeled at 77.5% and 77% efficiencies based on the relative absorbances at 280 nm and 650 nm.

### Preparation of mRNAs

The native mRNA rpl41a was generated by polymerase chain reaction-(PCR) linearizing a puc19-vector containing the rpl41a mRNA sequence downstream of T7 promoter followed by T7 *in vitro* transcription. The reverse primer for PCR encoded a poly(A)_30_ tail through a poly(dT)_28_ track and two ultimate 2’O-methylated uridine bases. Additionally, as a consequence of using a T7 promoter, two G nucleotides were appended to the 5’ end of the sequence to facilitate transcription. Thus, the final rpl41a sequence consisted of two G residues followed by the native rpl41a mRNA sequence and a 30-nucleotide poly(A) tail.

The 24mer used in Extended Data Figure 2. was generated by PCR linearizing and transcribing the first 45-nucleotides of our rpl41a mRNA construct. This 45mer was then hybridized to a 21-nucleotide DNA fragment containing a 5’ biotin, thus leaving the first 24-nuceotides—the 5’ UTR of rpl41a—single stranded.

All transcribed mRNAs were gel-purified using either 7% (rpl41a) or 12% (45mer) 19:1 acrylamide, 7M urea-tris, boric acid, EDTA (TBE) polyacrylamide gels. The mRNAs were capped using the vaccinia virus capping enzyme kit (New England Biolabs) following the manufacturer’s protocols. However, the capping reactions were allowed to proceed for 2 hours rather than 30 minutes. Capping reactions were then purified using Monarch RNA clean-up columns (New England Biolabs).

### Sequences of mRNA and oligonucleotides

rpl41a mRNA: GGAGACCACAUCGAUUCAAUCGAAAUGAGAGCCAAGUGGAGAAAGAAGAGAACUAGAAGACUUAAGAGAAAGAGACGGAAGGUGAGAGCCAGAUCCAAAUAAGCGGAUUAUGAGUAAAUAACUCUAAUUUUGUUUUAAAUUCUUUCAAGAGUAUCGUAAUGUCAUUGAUGAAUUAACAUGUUAGUUUCUAUUCUACCUCAUAAUGGAUCUAAAUUGCAUACUAAUCUCACGGUGGGGUGUAAACCAUUGCCUACUAUUUAUAUAGUGCUUUAUAUAUGUCUCACAUAGUUUAAUCAAUUGUCCGUUUUUUUGAAAAAAAAAAAAAAAAAAAAAAAAAAAAAA

rpl41a 24mer (bases hybridized to DNA oligo underlined): GGAGACCACATCGATTCAATCGAA<u>ATGAGAGCCAAGTGGAGAAAG</u>

rpl41a 51mer (Integrated DNA Technologies) GGAGACCACATCGATTCAATCGAAATGAGAGCCAAGTGGAGAAAGAAGAGA-FAM

DNA oligo for tethering rpl41a (Integrated DNA Technologies) Biotin-ACATGTTAATTCATCAATGAC

DNA oligo for tethering 24mer (Integrated DNA Technologies) Biotin-CTTTCTCCACTTGGCTCTCAT

Targeting Oligo for labeling rpl41a at position 14: CCGTCGCCATCTCCCGTAGGTGAAGGGCTTGAAGGTTCCATTCCCGATGTGGTCTC

Targeting Oligo for labeling rpl41a at position 92: CCGTCGCCATCTCCCGTAGGTGAAGGGCTTGGAGGTTCCATTCCCTGGCTCTCACC

### Fluorophore labeling of mRNAs

We utilized a terbium-assisted deoxyribozyme to generate fluorophore-labeled mRNAs as previously described^43^. Briefly, in a 500 µL total reaction volume, 10 µM rpl41a mRNA was hybridized with 12 µM targeting oligo and 17 µM ribozyme RNA component (rD) by heating to 65°C for 5 minutes and slow cooling to room temperature over an hour in 50 mM HEPES pH=7.4, 150mM sodium chloride (NaCl), and 2mM KCl. After hybridization, magnesium acetate (Mg(OAc)_2_) was added to 25 mM followed by the addition of 100 µM terbium (iii) chloride and 200 µM ethylenediamine-guanosine-5’-triphosphate (EDA-GTP). The reaction was allowed to proceed for 1 hour in a 37°C water path followed by 3 hours at room temperature, and then quenched by addition of 1/10^th^ volume 3 M sodium acetate pH=5.2 and ethanol precipitation. The ethanol precipitate was then resuspended and purified on a denaturing gel as described in the section above.

After purification of the site-specific EDA-GTP-mRNA conjugate, the mRNA was labeled using sulfo-Cy3-N-hydroxysuccinimide (Cy3-NHS). To perform fluorophore labeling, the mRNA was adjusted to 20 µM in 200 mM phosphate pH=8.4 containing 20% dimethylsulfoxide (DMSO) and 1 mM Cy3-NHS. The reaction was allowed to proceed for 45 minutes at room temperature, and then quenched with the addition of 1/10^th^ volume of 3 M sodium acetate pH=5.2. After quenching, excess dye was removed and the phosphate was desalted into 0.3 M sodium acetate using a 5 mL desalting column (Cytiva) and the eluate was ethanol precipitated. The purified, fluorescently labeled RNA was then capped following the procedure above.

### Assay buffer

All measurements were performed in a standard Reaction Buffer (20mM HEPES pH=7.4, 100mM KOAc, 3mM Mg(OAc)_2_) previously determined to facilitate *in vitro* translation initiation^30,35,40^.

### mRNA recruitment gel-shift assays

mRNA recruitment gel-shift assays used to monitor a stalled 48S-PIC on the start codon of rpl41a were performed using previously described standard protocols^11,30,35^. Briefly, *in vitro* transcribed rpl41a mRNA was ‘hot capped’ using a 1:100 ratio of α-P^32^-GTP:GTP. Mixtures containing the relevant initiation factor mixes (**Extended Data Figure 1b**), initiator tRNA, ATP•Mg^2+^, and 40S ribosomal subunits were assembled, and reactions were started with the addition of hot-capped mRNA. After half an hour, reactions were quenched by loading into a running a native 4% 37.5:1 acrylamide tris, HEPES, EDTA, magnesium (THEM) gel, which was then imaged on a Typhoon 5 (Cytiva) after exposure to a phosphor screen.

### Fluorescence anisotropy

For the titration-regime fluorescence anisotropy assays, experiments were performed using a 3’-fluoroscein-modified RNA fragment (Integrated DNA Technologies) consisting of the first 51 nucleotides of our rpl41a construct (51mer-FAM). Two-hundred microliter reactions were assembled by diluting a 100 µM of 51mer-FAM to 500 nM or 50 nM in reaction buffer. Two-hundred microliters was transferred to a quartz cuvette and small volumes of eIF4G were titrated into the cuvette to generate RNA:eIF4G stoichiometries of 0, 0.2, 0.4, 0.6, 0.8, 1, 1.5, 2, 2.5, 3, 3.5, 4, and 5 or 0, 0.2, 0.4, 0.5, 0.6, 0.8, 1, 1.2, 1.6, 2, 2.5, 3, and 4 for the 500 nM and 50 nM reactions respectively. The total titration did not change the total reaction volume by more than 10%. After each titration point, the sample was allowed to equilibrate for 5 minutes, and the fluorescence anisotropy^58^ was then measured using vertically polarized excitation light on a Horiba fluorimeter with excitation at 480 nm and emission at 518 nm.

### Total Internal Fluorescence (TIRF) microscopy

All smFRET experiments were performed on a lab-built, prism-type TIRF microscope described previously^59,60^. Briefly, samples were excited with a 532 nm laser (gem 532; Laser Quantum) at a power corresponding to 30 mW just before hitting the prism and imaged using an Andor Ultra 888 electron-multiplying charge-coupled device (EMCCD) camera. Movies were collected using an exposure time of 100 ms and were conducted with either continuous excitation by the laser or by mechanically shuttering the laser for 1 frame every 3 seconds.

All experiments were performed in quartz microfluidic flowcells passivated with a monolayer of a mixture of methoxy-terminated PEG and biotin-terminated PEG (5000 kDa, Laysan), as previously described^59,60^. In order to improve the previously described passivation, we altered the standard PEG-deposition steps to be a one-step passivation by directly using PEG-silane in cloud-point conditions. The final passivation solution was the bottom layer of a 5% PEG cloud-point solution generated in 1% acetic acid and 750 mM MgSO_4_, which was allowed to react with the quartz surface in a humidified chamber overnight. The ratio of methoxy-capped PEG to biotin-PEG was 10,000:1.

For all experiments, reaction chambers were prepared by washing the flowcells with Reaction Buffer, followed by Reaction Buffer containing 0.1% tween-20. Tween-20 was then expelled from the flowcell using Reaction Buffer and 10 nM streptavidin was allowed to bind the biotin-PEG for 1 minute before being washed out with Reaction Buffer. RNA molecules were then specifically tethered to the surface, through streptavidin, by hybridizing a short DNA oligo containing biotin to the middle of rpl41a. The mRNA was allowed to bind the surface for 30 seconds and then untethered mRNA molecules were expelled by washing the flowcell with Reaction Buffer. Finally, Imaging Buffer (Reaction Buffer supplemented with 1% glucose, 1 ug glucose oxidase, 0.85 <u>µ</u>g catalase, 1 mM cyclooctotatraene, 1 mM nitro-benzoic acid, 2 mM trolox aged to 2% oxidation in a 50 <u>µL volume</u>) containing triplet-state quenchers and an oxygen-scavenging system was injected into the flowcell prior to our injection experiments. For ejection experiments, this Imaging Buffer also contained whatever protein complex was ejected from the flowcell (discussed in detail in the next section).

### smFRET experiments

The association rates for Cy5-eIF4G/G:E were determined by injecting Cy5-eIF4G/G:E into flowcells containing capped, uncapped, or internal mRNAs at varying concentrations of Cy5-eIF4G/G:E. The concentration series consisted of 1 nM, 3 nM, and 5 nM. The association rates for Cy5-eIF4G:E+A/+AB were determined from a single concentration point where 3nM Cy5-eIF4G:E was mixed with either 2 µM eIF4A and 2 mM ATP or 2 µM eIF4A, 500 nM eIF4B and 2 mM ATP and subsequently injected into the flowcell.

Similarly, the pre-steady-state dissociation rates for Cy5-eIF4G/G:E were determined by allowing 3 nM Cy5-eIF4G/G:E to bind the surface-tethered mRNA for 5 minutes in Imaging Buffer and then ejecting all free, unbound Cy5-eIF4G/G:E from the flowcell by exchanging the solution in the flowcell with fresh imaging buffer which lacked Cy5-eIF4G:E. For experiments containing eIF4A and/or eIF4B, the experiment was performed by ejecting all of the free, unbound cy5-eIF4G/G:E using fresh Imaging Buffer containing 2 µM eIF4A and 2 mM ATP or 2 µM eIF4A, 500 nM eIF4B, and 2 mM ATP.

Cy5-eIF4E experiments were performed analogously to those of Cy5-eIF4G:E, except a titration series of 5 nM, 25 nM and 50 nM were used to determine the association rate of Cy5-eIF4E binding to each mRNA construct. Since binding was transient, no pre-steady-state experiments for Cy5-eIF4E were performed, but rather the steady-state dissociation rate was used. eIF4G:E-Cy5 injection and ejection experiments were performed at 5 nM to compensate for the lower labeling efficiency of eIF4E-Cy5.

For all non-shuttering injection and ejection experiments, the exchange of solution in the flowcell was started approximately 1.5 seconds (15 frames) after imaging began. For all shuttering experiments, ejection occurred between 3 and 6 seconds (frames 2 and 3).

### Analysis of smFRET data

All analyses of smFRET data were performed using custom written, in-lab, but publicly available software. Individual trajectories were extracted from movies using vbscope^61^, and trajectories were analyzed and fit to hidden Markov models using tMAVEN^62^, a Python-based software package which integrates single molecule analysis methodologies our laboratory has previously published into one software.

For each trajectory, E_FRET_ was calculated as the acceptor intensity divided by the sum of the donor and acceptor intensities. Then, the signal to background ratio was calculated for each trajectory, and trajectories with less than a 5 signal-to-background ratio were removed prior to analysis. The high signal-to-background ratio trajectories were then visually inspected to look for the characteristics of a single molecule—single-step photobleaching or single step photoblinking. The photobleaching point was then set by the user in tMAVEN^62^. Not all trajectories taken under shuttering conditions photobleached during the time course of the experiment. In this case molecules were selected that displayed similar photophysical and intensity profiles to the non-shuttering and shuttering trajectories which did photobleach, and the entire trajectory was utilized. After identification of single molecules and photobleaching or photoblinking, each E_FRET_ trajectory was idealized using a 3-state hidden Markov model using the vbFRET algorithm^63^. The Viterbi paths of each trajectory were then clustered using a k-means algorithm and reproducibly returned overall E_FRET_ values near 0.05, 0.35, and 0.7 for the 3 states for Cy5-eIF4G:E. These three E_FRET_ states were then simplified into a two-state ‘bound’ and ‘unbound’ model by combining the two higher E_FRET_ states into one ‘bound’ state and leaving the low E_FRET_ state representative of ‘unbound’ mRNA molecules. Notably, this is equivalent to running the 3-state HMM model and then thresholding at an E_FRET_ of 0.25 to separate the bound and unbound states. These final, idealized trajectories were then used to calculate the rates in each experiment.

Rates were calculated using survival analysis of the dwell time distributions in each experiment. For injection experiments, the pre-steady-state association rate was calculated by counting the dwell time points between injection and the first binding event (transition from the low E_FRET_ state to the combined high E_FRET_ state). These dwell time points were then converted to survival probabilities by calculating what fraction of the dwell time points were longer than time, t, for all possible t. The resultant survival probabilities were then fit with a monoexponential decay function (Equation 1). The rates determined from these exponential fits were then scaled linearly to account for incomplete labeling of our proteins by multiplying the rate determined from the fit by the inverse of the labeling efficiency. Notably, the labeling efficiency of Cy5-eIF4G and Cy5-eIF4G:E, from which most of the conclusions in this work are drawn, are nearly identical, so this correction factor primarily affects the comparison of Cy5-eIF4G:E and eIF4G:E-Cy5.

Similarly, the steady-state dissociation rates were calculated by counting the dwell time points in the bound and unbound states across every transition in the injection experiment, except for the first and last dwells, which correspond to the pre-steady-state event and photobleaching, respectively. These dwell time points were then converted to survival probabilities and fit to biexponential decay functions (Equation 2), because the process reports on both dissociation and facilitated dissociation (**Extended Data Figure 2a**). The steady-state rates are population weighted averages of the two phases of the biexponential fit (Equation 3). Consistent with existing literature^16,20,25^, all reported rates for Cy5-eIF4E in the absence of eIF4G are steady-state rates, and the reported association rates are the population-weighted average of a biexponential fit.

Similar to the pre-steady-state association rate, the pre-steady-state dissociation rate was calculated by counting all of the dwell times between the ejection event and the first dissociation event (transition from the combined high E_FRET_ state into the low E_FRET_ state). These dwell times were used to calculate survival probabilities which were fit with a monoexponential decay function (Equation 1).

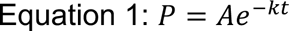

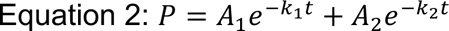

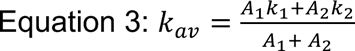

### False coloring of supplemental movies

The supplemental movies accompanying this article were false colored by manually contrasting the 16-bit gray scale raw images to an upper ceiling of ∼3000 and then down sampling to an 8-bit RGB movie in ImageJ^64^. Each channel was then manually false colored in ImageJ^64^ by adjusting the color balance for the red or green channel and sliding the respective color channel ceiling to a value of 127. This has the effect of giving each supplemental movie a similar level of brightness, although the movies are included to draw attention to the appearance and disappearance of spots during the depicted ejection/injection experiments, and the relative brightness of the spots are not relevant to the conclusions of the manuscript.

### Statistical analyses

The titration-regime fluorescence anisotropy experiments were performed once for each concentration.

All single-molecule experiments were performed as two biological replicates and nearly every individual single-molecule experiment contained >100 trajectories from which survival probabilities could be determined. However, experiments with low single-to-noise or trajectory numbers were repeated 3 times: These experiments were: ejection of Cy5-eIF4G:E from the internal mRNA, and ejection of Cy5-eIF4G:E from the internal mRNA using buffer supplemented with eIF4A and ATP. The titration-regime anisotropy experiments were performed once at each concentration of RNA.

The association rates for Cy5-eIF4G/G:E and Cy5-eIF4E with the mRNA (Figures 1 and 2) were determined by fitting the concentration-dependent association rates from individual experiments in the concentration series to a line. The reported rates for those experiments are the slopes of the lines and the errors are errors of the fit. All other reported rates are the means of the replicates, and the errors are the standard deviations of those replicates. The raw data points used to determine the means and standard deviations are plotted on each bar plot.

**Figure 2.**
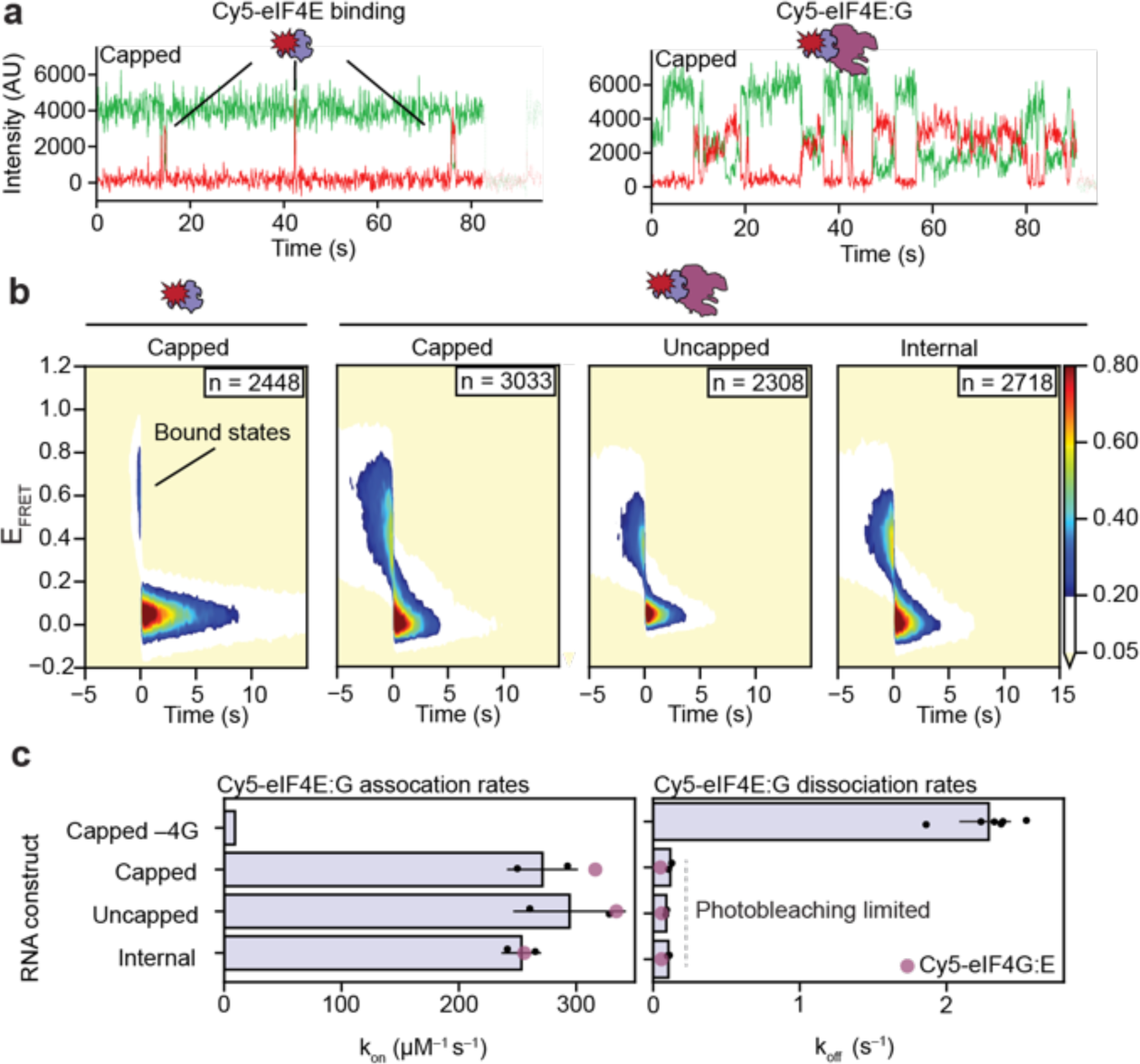
eIF4E functions in concert with eIF4G irrespective of the cap. (a) Example trajectories of Cy5-eIF4E binding to the capped mRNA construct in the absence (left) or presence (right) of unlabeled eIF4G. In the absence of eIF4G, Cy5-eIF4E only transiently samples the cap, whereas it stably binds in the presence of eIF4G. (b) Surface contour plots post-synchronized to the steady-state transitions of eIF4E out of the bound state: Time = 0 marks the time at which eIF4E dissociates from the vicinity of the donor fluorophore, such that the negative time axis indicates how long Cy5-eIF4E remains bound before dissociation and the positive time axis shows the delay time between binding events. (c) Quantitation of the association and dissociation rates of Cy5-eIF4E and eIF4G:E-Cy5 from the three mRNA constructs. The association rates for the capped, uncapped, and internal constructs are determined from experiments performed at 5nm eIF4G:E-Cy5, whereas the capped–4G association rate is derived from the slope of the apparent association rates of Cy5-eIF4E performed across three concentrations. The nearly identical rates derived from equivalent experiments performed with Cy5-eIF4G, plotted in Figure 1, are shown as purple dots to aid the comparison between Cy5-eIF4E and Cy5-eIF4G.

**Figure 3.**
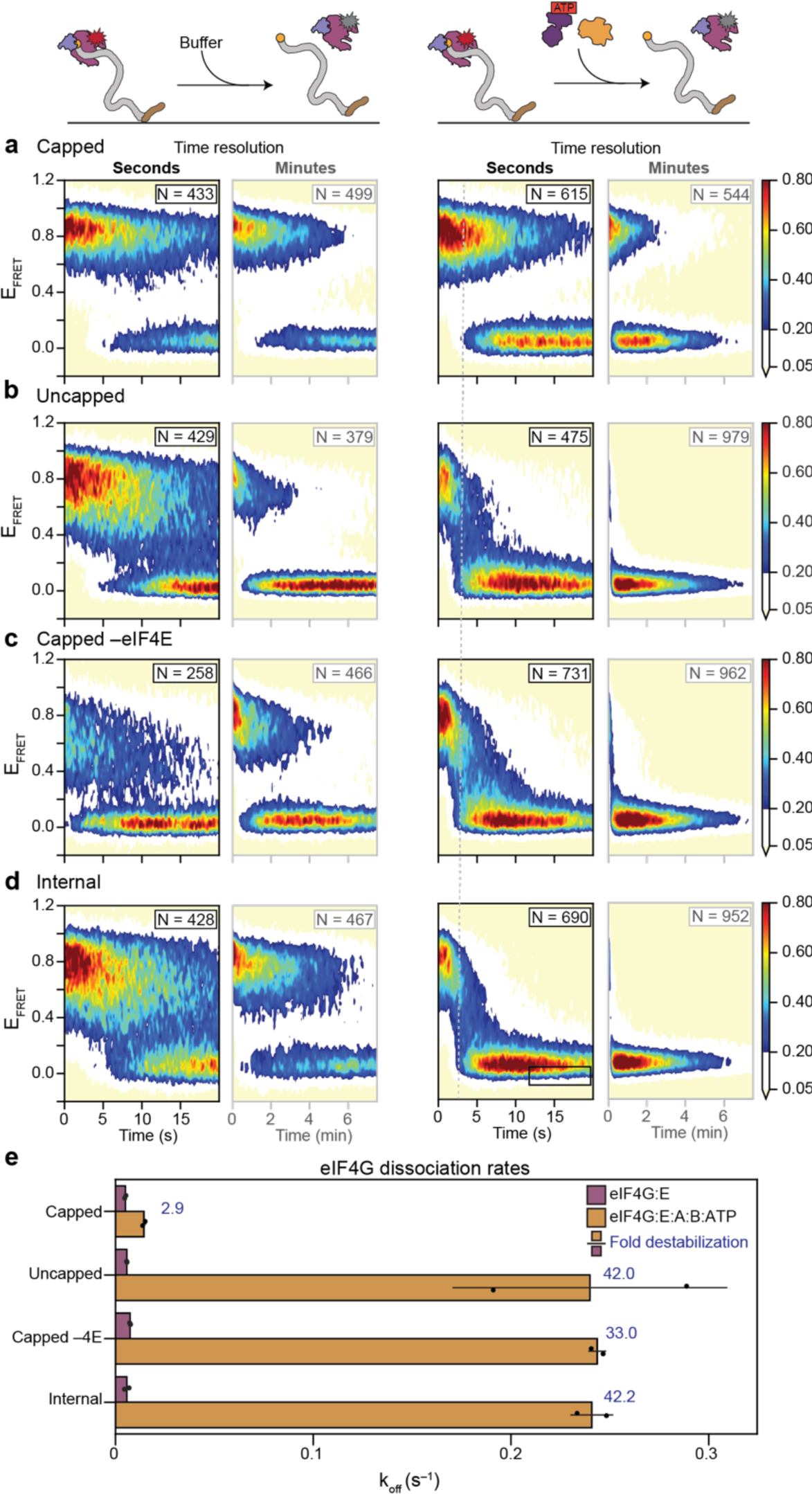
eIF4A strips eIF4G:E from RNA. Surface contour plots showing ejection experiments where excess, unbound Cy5-eIF4G:E in the flowcell is replaced with buffer lacking Cy5-eIF4G:E or replaced with buffer lacking Cy5-eIF4G:E but supplemented with eIF4A, eIF4B, and ATP (right). Each experiment was recorded at both 10 frames per second (fps) and 1/3 fps to obtain kinetic information in both the seconds (black) and minutes (gray) timescales. These experiments were performed on the (a) capped, (b) uncapped, (c) capped RNA without eIF4E, and (d) internal mRNA constructs. Each experiment begins with Cy5-eIF4G:E bound to the RNA. E_FRET_ starts at a high value and remains stable until photobleaching in the absence of eIF4A, eIF4B and ATP. However, in the presence of eIF4A, eIF4B and ATP, EFRET rapidly decays within ∽3 seconds (guiding line). This indicates that eIF4A strips eIF4G:E from RNA. eIF4E recognition of the cap (a) is able to resist this stripping. (e) Pre-steady-state off rates were calculated from the individual trajectories of the experiments performed in (a)-(d). These rates show eIF4G is destabilized by 30-40 fold in the presence of eIF4A, eIF4B, and ATP, but eIF4G is resistant to this destabilization in the presence of both eIF4E and the cap.

**Figure 4.**
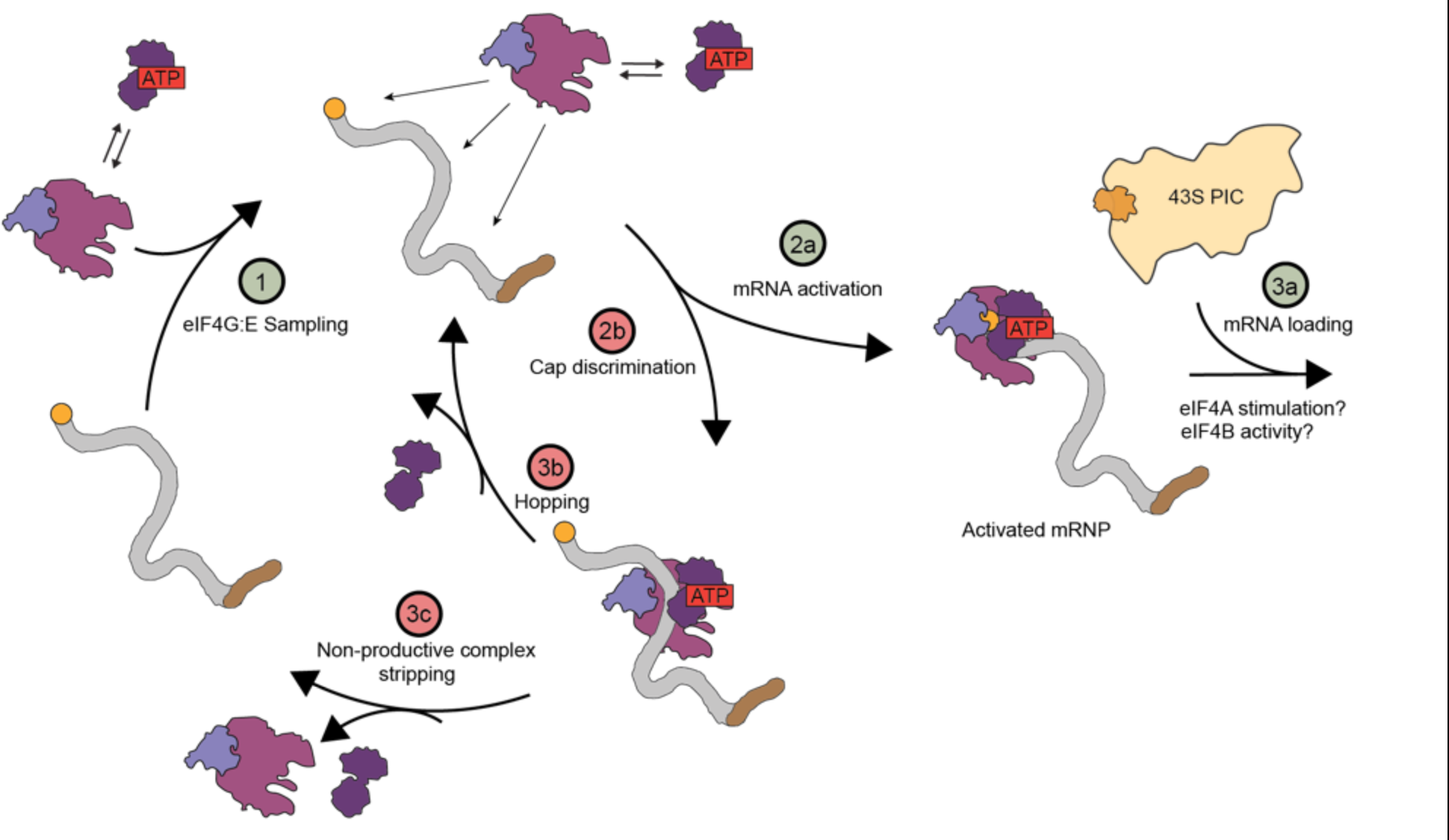
The mechanism of mRNA activation. Our experiments lead us to the following new model of mRNA activation: In the first step, eIF4G:E nonspecifically samples and binds to mRNA. At the second step, the pathway bifurcates into productive (green) or nonproductive (red) pathways. In the non-productive fork (2b), eIF4G:E binds far away from the cap, preventing eIF4E from associating with the cap, allowing eIF4A to recycle the eIF4F complex by either facilitating its intramolecular hopping to another place on the same mRNA molecule (3b) or completely removing it from the RNA (3c). In the productive fork, eIF4G:E binds at the 5’ UTR near the cap, and the eIF4E-cap interaction allosterically signals to eIF4A, through eIF4G, not to strip eIF4G. This stalled eIF4F complex at the cap constitutes the previously elusive activated mRNP, in which we speculate eIF4A may be held in an inactive conformation which can be reactivated by an eIF4B containing 43S-PIC prior to loading the mRNA onto the ribosome (3a). Notably, because eIF4A is present at high concentrations in the cell, this model is agnostic to which step eIF4A binds eIF4G:E, and due to its modest affinity, we speculate it is dynamically sampling the complex throughout the cycle.

The number of trajectories utilized, number of events in the fit, and all fitting parameters for the survival-analysis of every single-molecule experiment can be found in **Supplemental Table 1.**

## Acknowledgements

This work was supported by funds to R.L.G from the NIH (5R01GM084288 and CA277727). R.C.G and N.A.I were supported by NSF GRFP fellowships (DGE – 1644869). E.W.H. was funded through an F32 fellowship from the NIH (GM139360). Access to the Horiba Fluorimeter was enabled through NSF award 1828491. The authors would like to thank the Columbia Precision Biomolecular Characterization Facility for access to equipment.

## Author contributions

This work was conceptualized by R.L.G and R.C.G. The single molecule experiments were performed, analyzed, and interpreted by R.C.G with input from R.L.G and N.A.I.. The mRNA recruitment gel-shift experiments were performed, analyzed, and interpreted by N.A.I with input from R.L.G and R.C.G.. R.C.G and N.A.I. generated the materials with contributions from V.M.C and E.W.H. The improved surface passivation was conceptualized by C.K.T. and performed by R.C.G. R.L.G and R.C.G. wrote the paper.

## Data availability

All data analyzed in this study are available in the manuscript and Extended Data. The number of molecules included and the parameters for the raw survival curve fit for each replicate, uncorrected for framerate and labeling efficiency, are available in Supplemental Table 1. The large, raw movie files are available upon request to R.L.G.

## Code availability

The code used in this study is freely available at our group’s github page. Vbscope: https://github.com/GonzalezBiophysicsLab/vbscope-paper

tMAVEN: https://github.com/GonzalezBiophysicsLab/tmaven

---

**Extended Data Figure 1.**
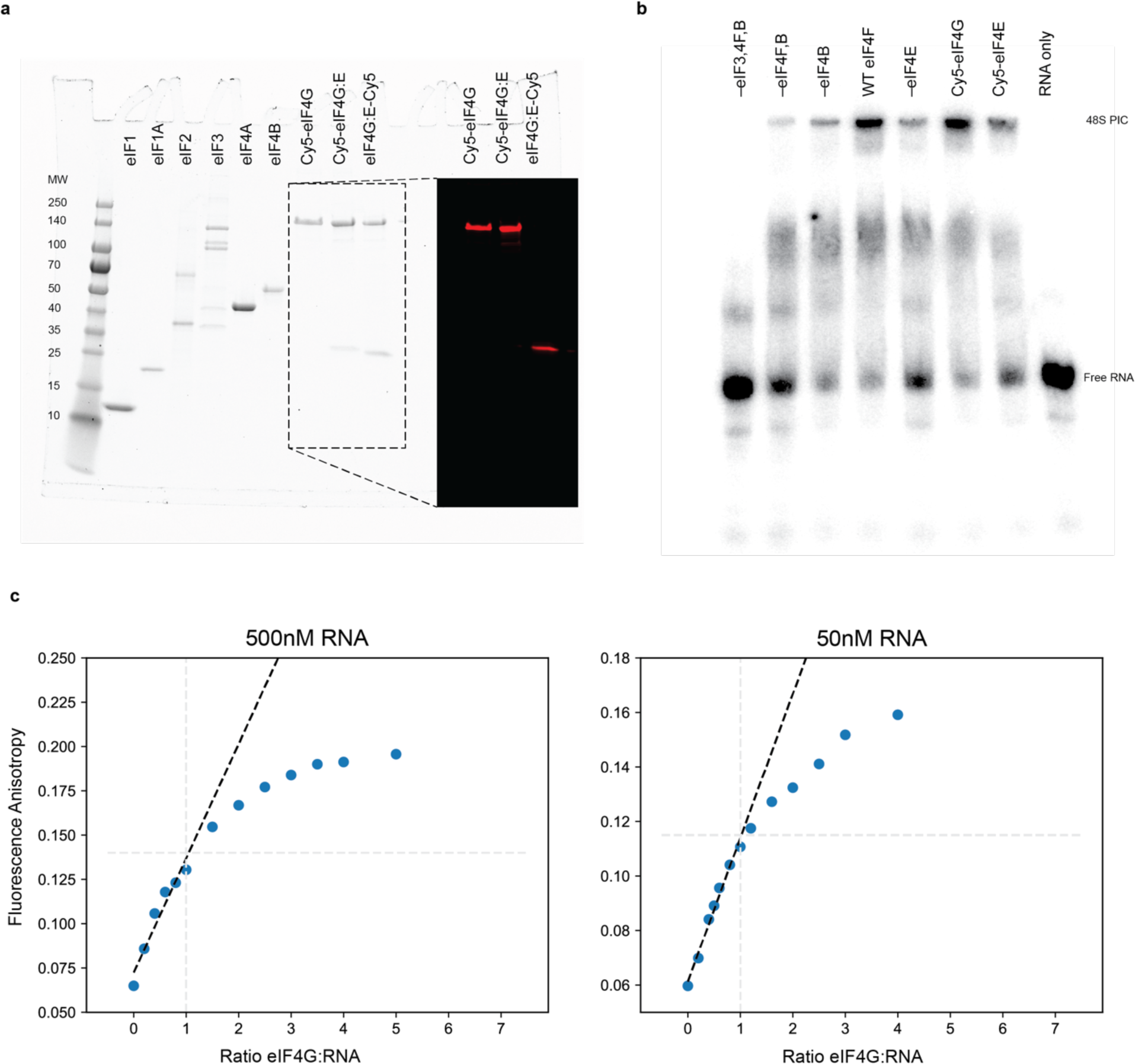
Purification, labeling, and *validation of biochemical components*. **(a)** SDS-PAGE gel showing all of the purified eIFs required to perform mRNA recruitment gel-shift assays. The inset shows a red fluorescence scan of the outlined portion of the SDS-PAGE gel demonstrating that the factors are fluorescently labeled. **(b)** Standard mRNA recruitment gel-shift assay showing recruitment of mRNA onto a stable 48S-PIC in the presence of eIF4F. As expected, diminished recruitment is observed in the absence of eIF4B or eIF4E. **(c)** Titration-regime, anisotropy-based RNA binding experiments performed with either 500nM or 50nM FAM-labeled, 51-nucleotide rpl41a fragment. These experiments show that anisotropy continues to increase beyond a 1:1 molar ratio of mRNA:eIF4G, confirming multiple eIF4G molecules bind the short RNA, as previously observed. Each titration-regime experiment was performed once.

**Extended Data Figure 2.**
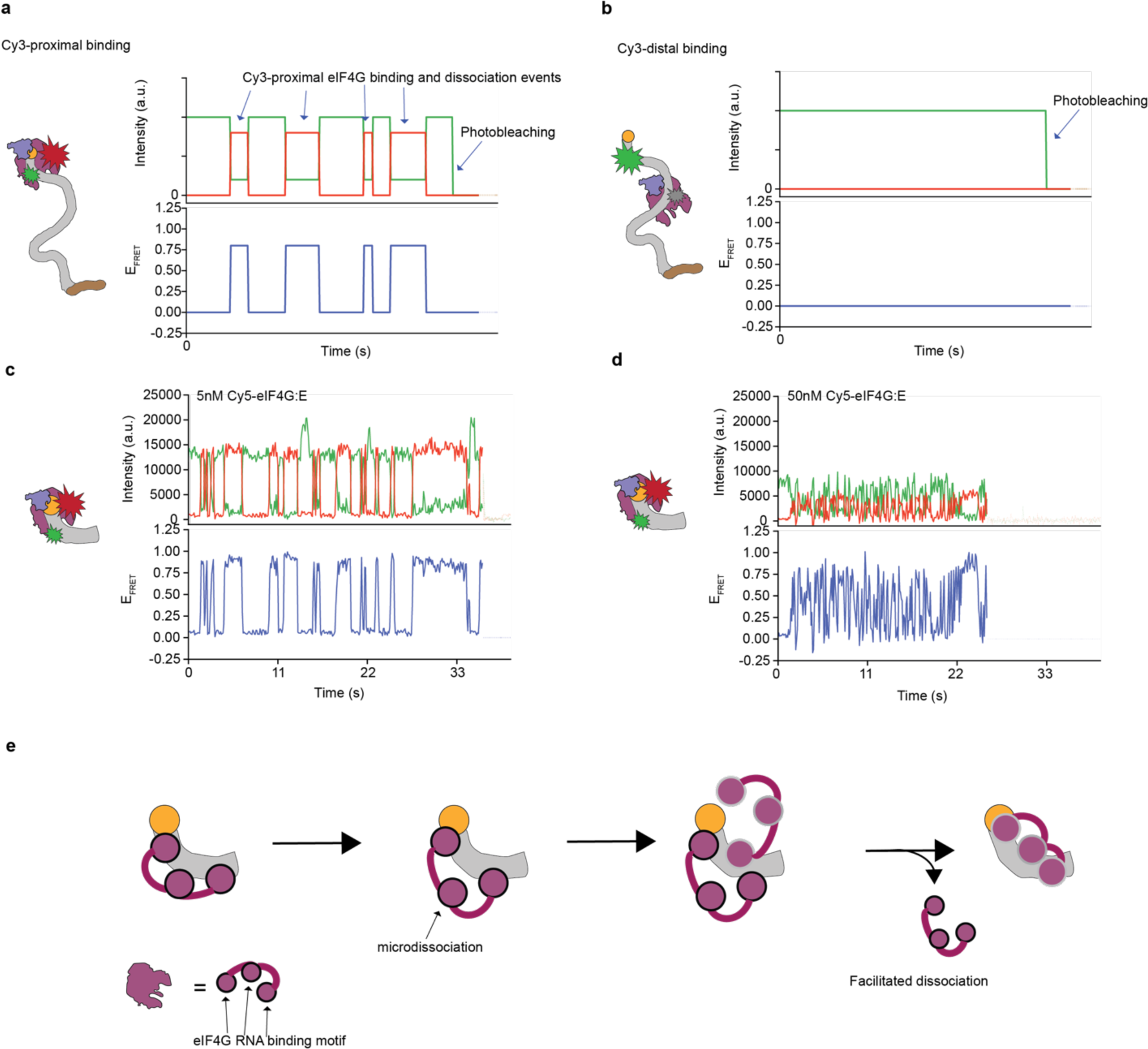
Validation of smFRET between Cy5-eIF4G:E and Cy3-mRNA. **(a)** Hypothetical intensities/FRET trajectories showing what is expected to happen if Cy5-eIF4G:E binds at a cap-proximal location on the capped mRNA construct . Upon binding, the green fluorescence intensity decreases while the red intensity increases for the duration of the binding event, leading to an increase in EFRET while Cy5-eIF4G:E is bound near the donor fluorophore. **(b)** Shows similar hypothetical trajectories, but in these trajectories Cy5-eIF4G:E binds at a cap-distal location far away from the donor fluorophore. Because the acceptor and donor fluorophores are not near each other, no changes in intensity or EFRET are detected before photobleaching. **(c)** Shows an example experimental trajectory for Cy5-eIF4G:E binding to the short, 24-nucleotide 5’ UTR of rpl41a. Because the RNA is short, we expect only a single binding site. At 5nM, we observe transient binding, but at 50nM **(d),** both the association and dissociation kinetics increase, indicating that eIF4G is able to facilitate its own dissociation. Facilitated dissociation is cartooned in **(e)** whereby the release of one of RNA binding domains of eIF4G, also called microdissociation, opens up a binding location for a new molecule of eIF4G, which then facilitates the release of the first eIF4G molecule.

**Extended Data Figure 3.**
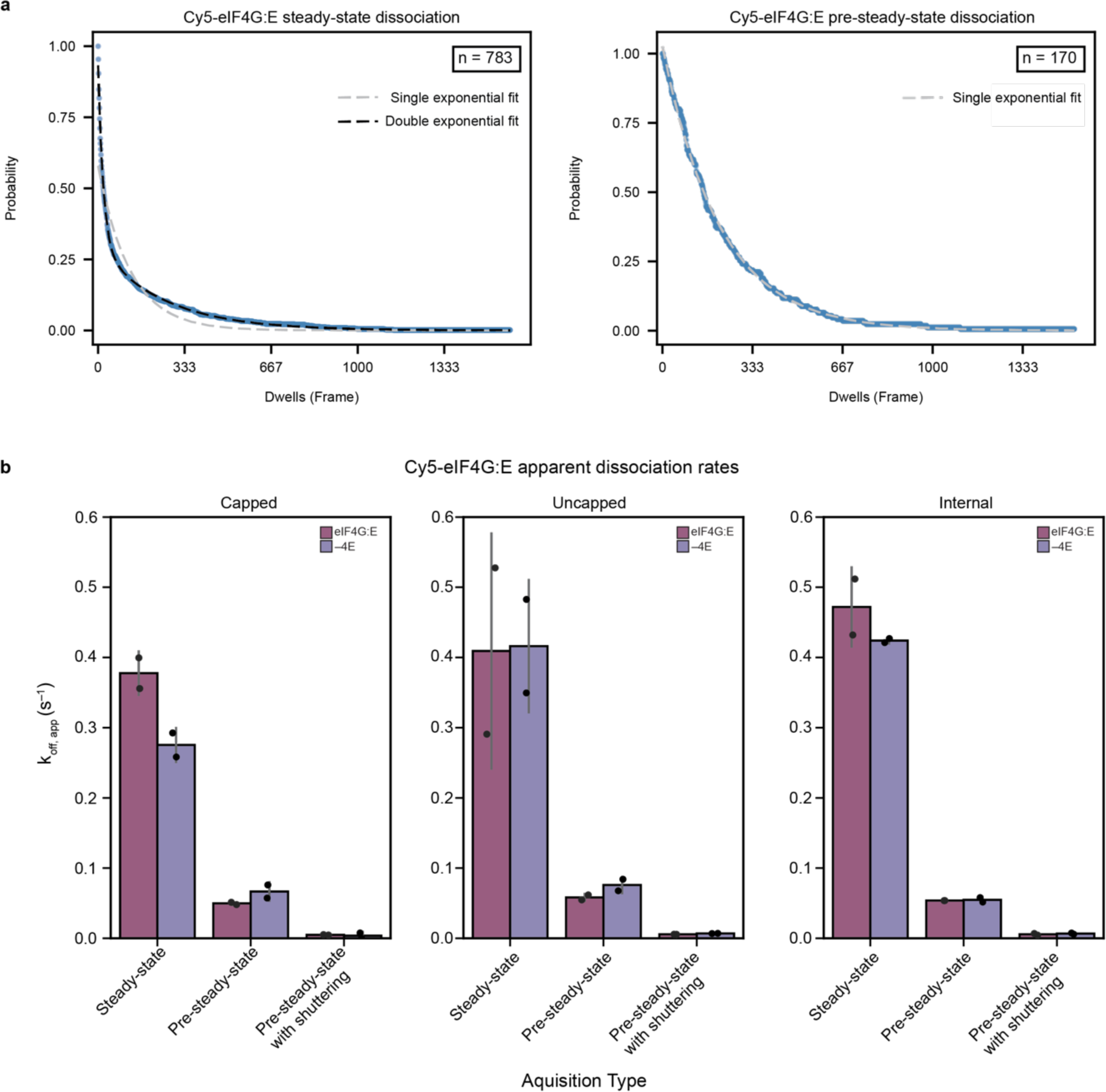
Cy5-eIF4G:E facilitates its own dissociation on full-length rpl41a mRNA. **(a)** Survival-time analysis showing that the steady-state dissociation (left) of Cy5-eIF4G:E on the capped mRNA construct is biphasic, whereas the pre-steady-state dissociation rate (right) is monophasic, indicating that steady-state dissociation events are a mixture of facilitated and non-facilitated dissociation events. **(b)** Comparison of the rate constants derived from steady-state and pre-steady-state experiments again demonstrate that eIF4G:E facilitates its own dissociation in the steady-state experiments. Thus, pre-steady-state experiments are more reflective of the inherent dissociation rate of eIF4G:E. However, comparison of the pre-steady-state dissociation rates recorded with and without shuttering of the excitation light source demonstrate that the pre-steady-state dissociation rate is limited by photobleaching. The steady-state dissociation rates plotted are population weighted averages of the two phases shown in **(a)**.

**Extended Data Figure 4.**
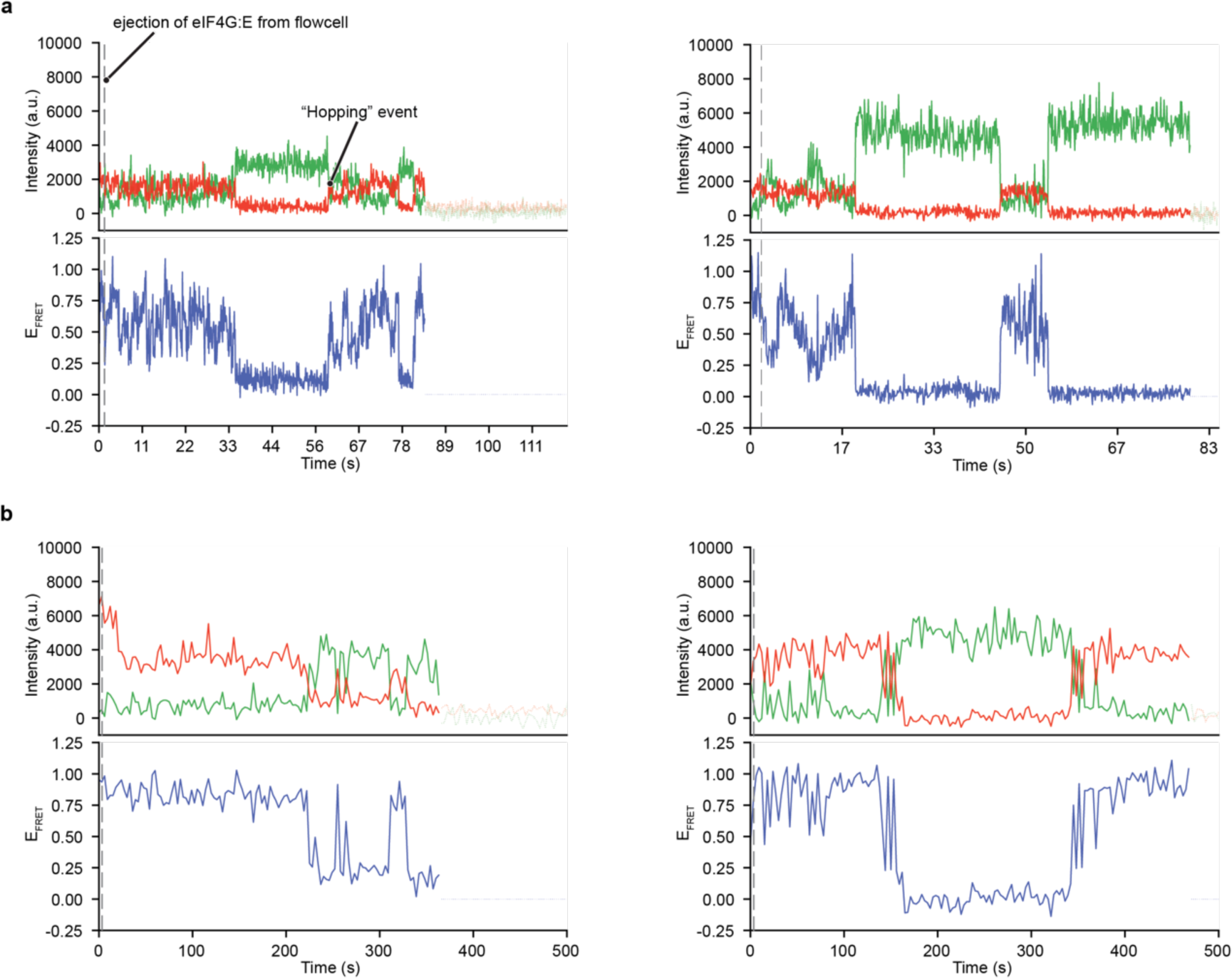
eIF4G:E can “hop” along an mRNA. Example trajectories taken from pre-steady-state ejection experiments on the capped construct where Cy5-eIF4G:E dissociates from the local position on the mRNA, causing EFRET to drop to 0, but later recovers, despite the all free eIF4G:E being removed from the flowcell, when the same, or another eIF4G:E bound elsewhere on the mRNA, microdissociates and then “hops” to the local position near the donor fluorphore on the mRNA. These example trajectories are shown for both continuous illumination **(a)** and excitation light shuttering **(b)** experiments.

**Extended Data Figure 5.**
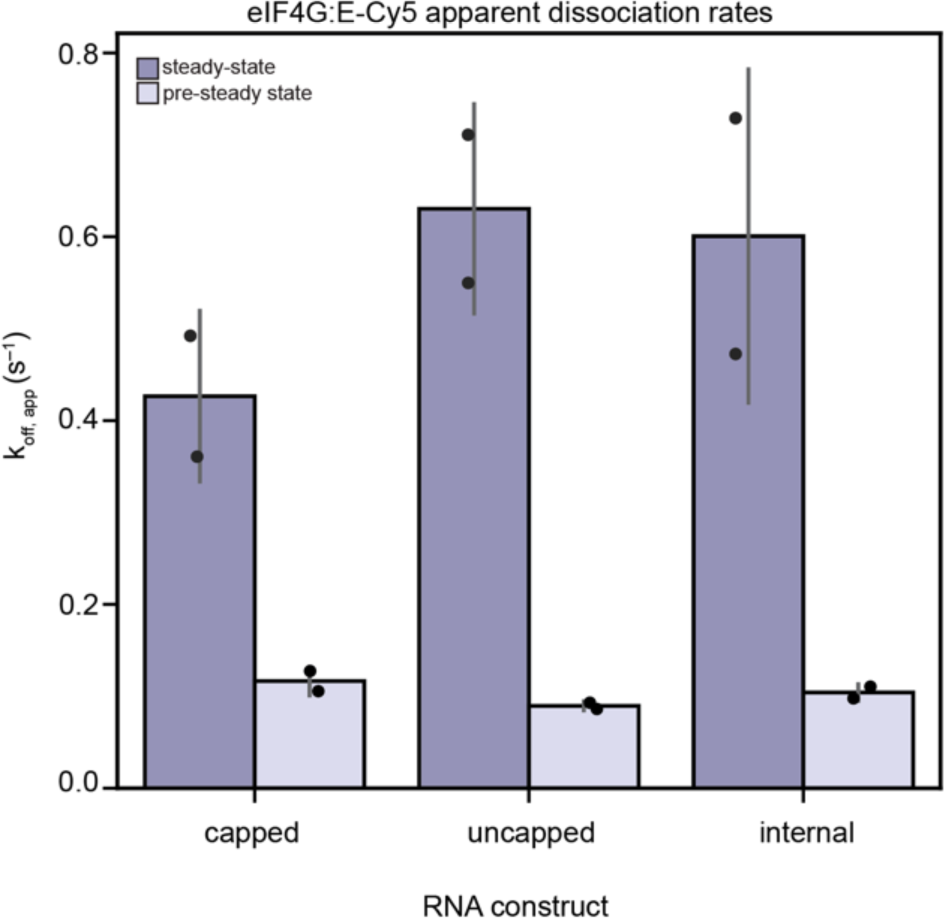
eIF4G:E-Cy5 facilitates its own dissociation. A bar graph showing the population-weighted average, steady-state dissociation rate of eIF4G:E compared to the pre-steady-state dissociation rate. Similar to Cy5-eIF4G/G:E, free, unbound, eIF4G:E-Cy5 is able to exchange with bound eIF4G:E-Cy5 leading to an increased apparent dissociation rate in steady-state experiments. This serves as additional evidence that eIF4G and eIF4E function as a complex.

**Extended Data Figure 6.**
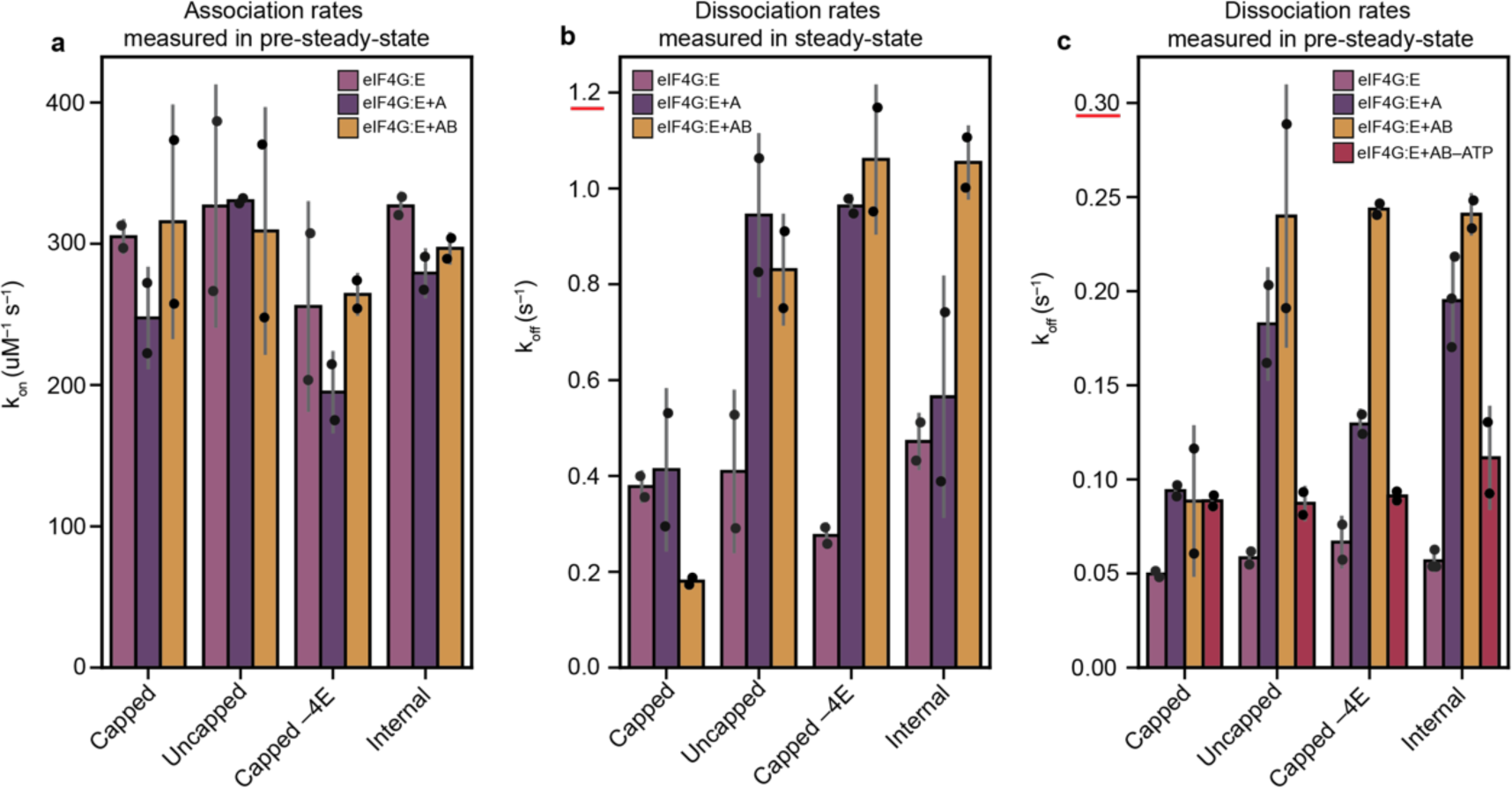
Association and dissociation kinetics of Cy5-eIF4G:E in the presence of eIF4A and eIF4B. **(a)** The pre-steady-state association rates calculated from a Cy5-eIF4G:E concentration series, also plotted in Figure 1, and the pre-steady-state association rates calculated from a single Cy5-eIF4G:E concentration point of 3 nM in the presence of saturating eIF4A+ATP or eIF4A+eIF4B+ATP. eIF4A and eIF4B do not appear to alter the association kinetics. **(b)** The steady-state dissociation kinetics calculated from the steady-state portion of the injection experiments for eIF4G:E, eIF4G:E+eIF4A+ATP, and eIF4G:E+eIF4A+eIF4B+ATP. eIF4A and eIF4B enhance the steady-state dissociation rate on every construct except the capped construct in the presence of eIF4E. **(c)** The equivalent pre-steady-state experiment for those described in **(b)**. eIF4A destabilizes eIF4G/G:E from each mRNA construct in an ATP-dependent manner except in the presence of both eIF4E and the cap. A red line at the highest tick mark on the Y-axis of **(b)** and **(c)** denotes the change in axis scale between the two graphs—highlighting that the steady-state dissociation rate far exceeds the pre-steady-state dissociation rate due to facilitated dissociation.

**Extended Data Figure 7.**
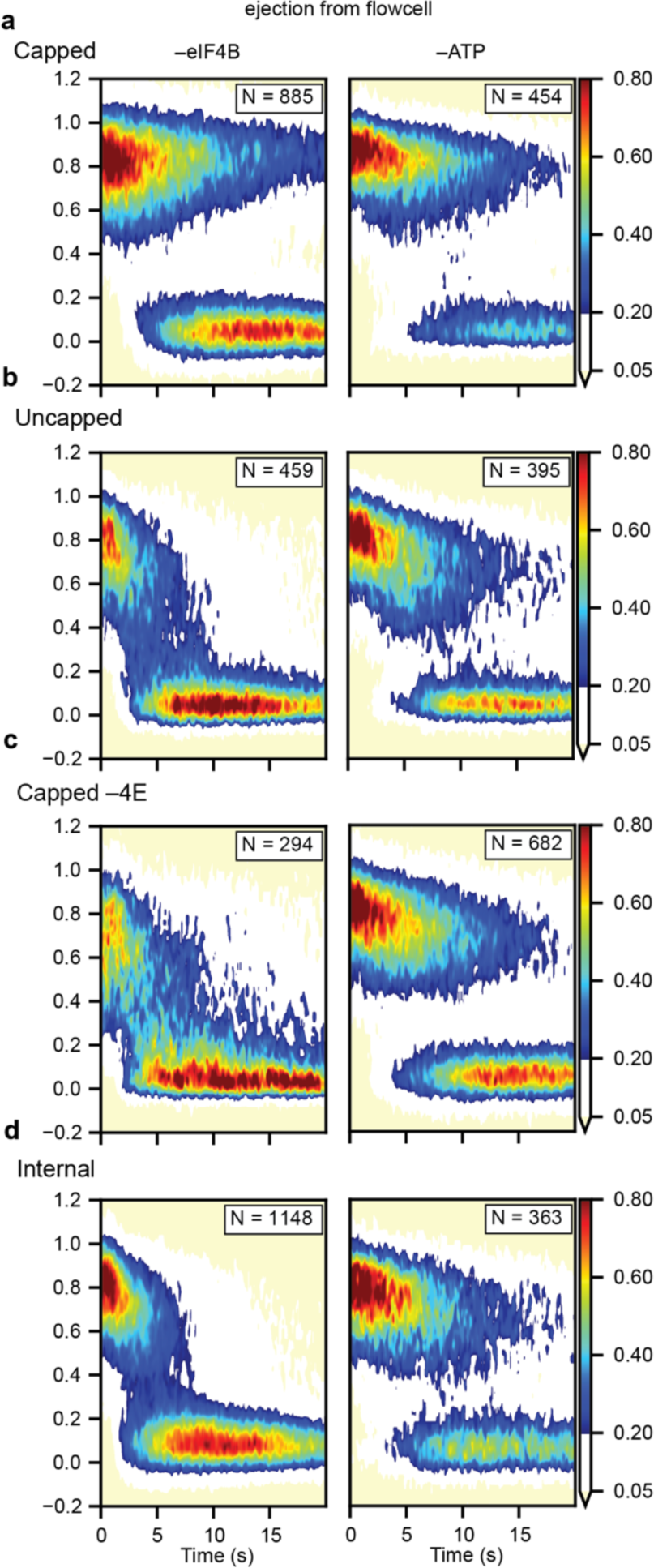
ATP, but not eIF4B, is required for stripping eIF4G from mRNA. Surface contour plots showing pre-steady-state ejection experiments. These ejection experiments were performed by exchanging all of the free, unbound Cy5-eIF4G:E in the flowcell with buffer lacking Cy5-eIF4G:E, but containing some combination eIF4A, eIF4B and ATP similar to Figure 4. On the left, reactions were performed without eIF4B, while on the right they were performed without ATP. All experiments were performed on the **(a)** capped, **(b)** uncapped, **(c)**, capped without eIF4E and **(d)** internal mRNA constructs. These results show efficient stripping of Cy5-eIF4G:E is not dependent on eIF4B, but is dependent on ATP.

**Extended Data Figure 8.**
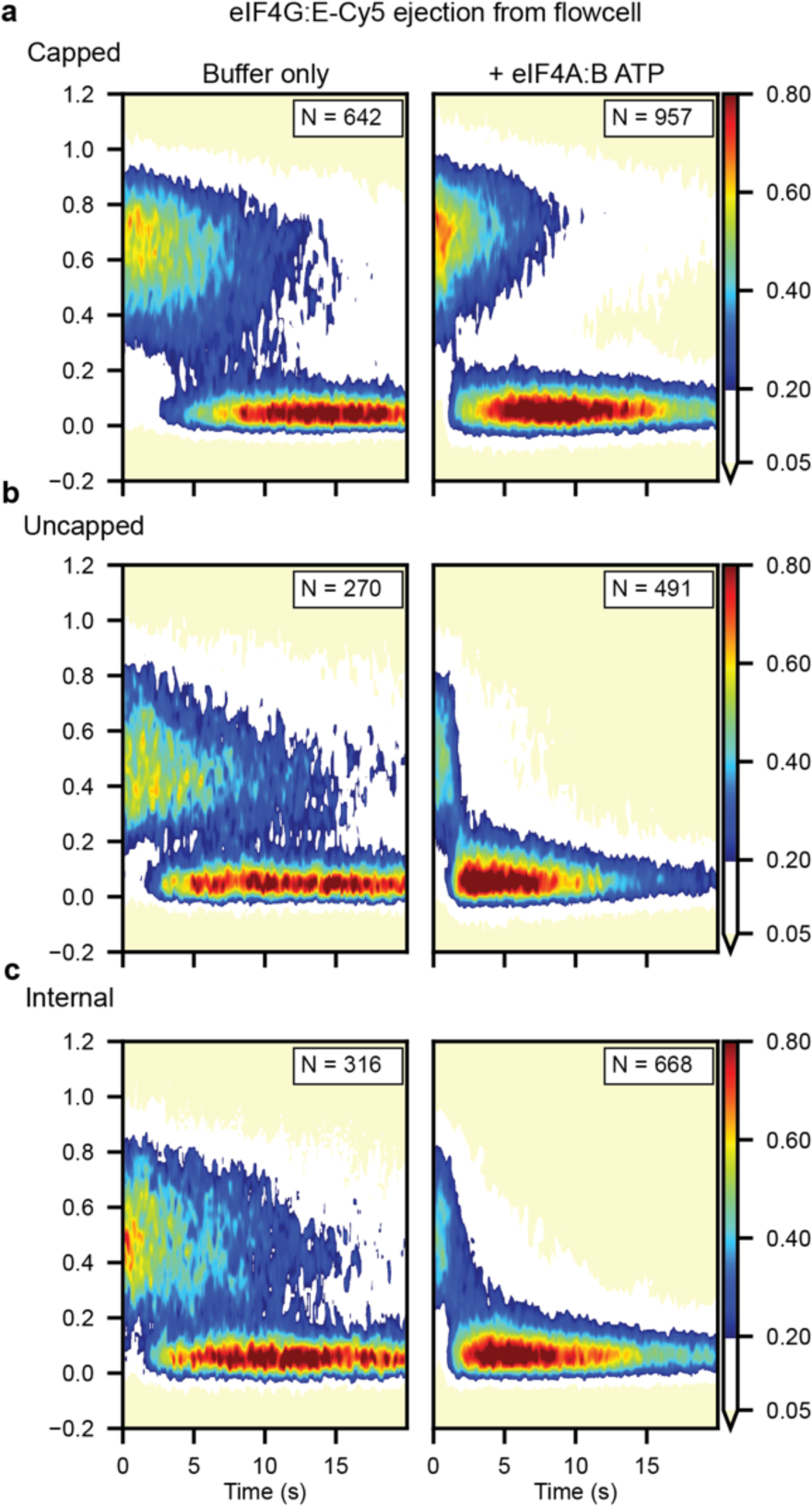
eIF4G:E is stripped from mRNA as a complex. The equivalent ejection experiments shown in Figure 4 but with the acceptor fluorophore on eIF4E instead of eIF4G. Similar to with Cy5-eIF4G:E, eIF4G:E-Cy5 is stripped from every position except from the cap. This demonstrates that eIF4G:E is stripped as a complex during the search for the cap.

**Extended Data Table 1.** Comparison of Cy5-eIF4G and Cy5-eIF4E results.

| Association rate constants measured in pre-steady-state, $k_{on}$ ($\mu M^{-1} S^{-1}$) | | |
| --- | --- | --- |
|  | Cy5-4G:E | 4G:E-Cy5 |
| mRNA construct |  |  |
| Capped -4E | 237 ± 41 | N.D. |
| Capped | 316 ± 5 | 271 ± 30<sup>a</sup> |
| Uncapped | 334 ± 5 | 294 ± 48<sup>a</sup> |
| Internal | 256 ± 39 | 253 ± 17<sup>a</sup> |

<sup>a</sup>rates calculated from single concentration point at 5nM

**Extended Data Table 1. Comparison of Cy5-eIF4G and Cy5-eIF4E results.**
| Dissociation rate constants measured in pre-steady-state, $k_{off}$ ($s^{-1}$) | | | | |
| --- | --- | --- | --- | --- |
|  | Cy5-4G:E | 4G:E-Cy5 | Cy5-4G:E:AB | 4G:E-Cy5:AB |
| mRNA construct |  |  |  |  |
| Capped -4E | 0.067 ± .013<sup>a</sup> | N.D. | 0.244 ± .004 | N.D. |
| Capped | 0.050 ± .002<sup>a</sup> | 0.116 ± .015<sup>a</sup> | 0.088 ± .039<sup>a</sup> | 0.165 ± .006<sup>a</sup> |
| Uncapped | 0.057 ± .005<sup>a</sup> | 0.090 ± .005<sup>a</sup> | 0.240 ± .069 | 0.590 ± .049 |
| Internal | 0.058 ± .005<sup>a</sup> | 0.104 ± .009<sup>a</sup> | 0.241 ± .011 | 0.373 ± .008 |

<sup>a</sup>Rates calculated from continuous acquisition, photobleaching limited

**Extended Data Table 2.** Summary of rate constants.

| Summary of rate constants |  |  |  |
| --- | --- | --- | --- |
| | $k_{on}$ ($\mu\text{M}^{-1}$) | $k_{off}$ ($\text{s}^{-1}$) | $K_D$ (pM) |
| Capped |  |  |  |
| eIF4G | $237 \pm 40^a$ | $0.0074 \pm 0.0003^a$ | $31 \pm 5$ |
| eIF4G:E | $316 \pm 5^a$ | $0.0050 \pm 0.0004^a$ | $16 \pm 1$ |
| eIF4G:E+A | $247 \pm 35^a$ | $0.0939 \pm 0.0042^{b,c}$ | $< 380 \pm 57^d$ |
| eIF4G:E+A+B | $315 \pm 81^a$ | $0.0144 \pm 0.0009^a$ | $46 \pm 12$ |
| eIF4G+A | $194 \pm 28^a$ | $0.1293 \pm 0.0073^b$ | $664 \pm 103$ |
| eIF4G+A+B | $264 \pm 14^a$ | $0.2436 \pm 0.0043^b$ | $922 \pm 52$ |
| Uncapped |  |  |  |
| eIF4G | $149 \pm 7^a$ | $0.0069 \pm 0.0002^a$ | $46 \pm 3$ |
| eIF4G:E | $334 \pm 5^a$ | $0.0057 \pm 0.0001^a$ | $17 \pm 1$ |
| eIF4G:E+A | $330 \pm 3^a$ | $0.1823 \pm 0.0293^b$ | $553 \pm 89$ |
| eIF4G:E+A+B | $309 \pm 87^a$ | $0.2399 \pm 0.0691^b$ | $776 \pm 312$ |
| Internal |  |  |  |
| eIF4G | $277 \pm 33^a$ | $0.0066 \pm 0.0012^a$ | $24 \pm 5$ |
| eIF4G:E | $256 \pm 39^a$ | $0.0057 \pm 0.0016^a$ | $22 \pm 7$ |
| eIF4G:E+A | $279 \pm 17^a$ | $0.1949 \pm 0.0241^b$ | $698 \pm 96$ |
| eIF4G:E+A+B | $297 \pm 10^a$ | $0.2408 \pm 0.0105^b$ | $812 \pm 45$ |

<sup>a</sup>Rates calculated from movies with framerate of 10 frames per second (fps) <sup>b</sup>Rates calculated from movies with framerate of 1/3 fps <sup>c</sup>Photobleaching limited <sup>d</sup> $K_D$ is the upperbound due to photobleaching limited off rate

