## Supplementary material for "The mechanism of mRNA activation": SI_Legends

**Supplementary Table 1. Single-molecule rates.** Supplementary table 1 contains the raw uncorrected rates, number of trajectories, and survival fitting parameters for the pre-steady-state and steady-state portions of every experiment reported in the article. The rates reported are all uncorrected for framerate, and therefore represent the rate *per frame* rather than s^–1­^. Additionally, the on rates have not been corrected for labeling efficiency. However, the framerates used to collect the data are also included in the table, and the materials and methods contains the labeling efficiencies. The different experiment types are separated into tabs eg eIF4G-Cy5 injection and eIF4G-Cy5 ejection.

**Supplementary Movie 1.** **Cy5**-**eIF4G:E rapidly associates with the 24mer.**

This injection experiment shows rapid association of Cy5-eIF4G:E with the 24-nucleotide 5’ UTR of rpl41a. Upon binding around ~1.2 seconds, the spots in the green channel disappear and corresponding spots in the red channel appear. This corresponds to FRET between the Cy3 donor on the mRNA and Cy5 acceptor on eIF4G.

**Supplementary Movie 2. Cy5-eIF4G:E rapidly associates with capped-rpl41a.**

This injection experiment shows the rapid association of Cy5-eIF4G:E with capped-rpl41a mRNA near the cap-proximal fluorophore. Upon binding around ~1.2 seconds, the spots in the green channel disappear and corresponding spots in the red channel appear. This corresponds to FRET between the Cy3 donor on the mRNA and Cy5 acceptor on eIF4G

**Supplementary Movie 3. Cy5-eIF4G:E rapidly associates with uncapped-rpl41a.** This injection experiment shows the rapid association of Cy5-eIF4G:E with uncapped-rpl41a mRNA near the cap-proximal fluorophore. This movie is indistinguishable from Supplementary Movie 2, demonstrating that the cap does not affect eIF4G:E association with the mRNA 5’ end. Upon binding around ~1.2 seconds, the spots in the green channel disappear and corresponding spots in the red channel appear. This corresponds to FRET between the Cy3 donor on the mRNA and Cy5 acceptor on eIF4G.

**Supplementary Movie 4.** **Cy5-eIF4G:E stably associates with mRNA.**

In this ejection experiment, 3nM Cy5-eIF4G:E was allowed to bind capped-rpl41a near the cap-proximal fluorophore. Between frames 2 and 3, all excess, unbound Cy5-eIF4G:E was removed from the flowcell and the bound Cy5-eIF4G:E was observed for dissociation. The excitation light was shuttered every 30 seconds to preserve the fluorophore. The spots in the red channel, corresponding to bound eIF4G:E, survive for upwards of tens of minutes—until RNAse is injected into the flowcell at 25 minutes. This illustrates just how stably eIF4G:E remains bound to mRNA.

**Supplementary Movie 5. eIF4E-Cy5 weakly associates with mRNA in the absence of eIF4G.** This pair of injection experiments demonstrate that in the absence of eIF4G, eIF4E only transiently samples the cap. On the left, Cy5-eIF4E dynamically samples the cap of capped-rpl41a with a cap-proximal dye, resulting in very short blips of red intensity in the red channel. In contrast, when co-injected with equimolar amounts of unlabeled eIF4G (right), Cy5-eIF4E rapidly and stably associates with the mRNA in a manner identical to that shown for Cy5-eIF4G:E.

**Supplementary Movie 6. eIF4G:E-Cy5 association with mRNA is not cap specific.** Although eIF4E binding to mRNA is known to be specific for the cap, co-injection of eIF4G and Cy5-eIF4E results in rapid and stable binding of the eIF4G:E-Cy5 complex to both uncapped mRNAs (left) and internal positions on the mRNA (right). This demonstrates that eIF4E non-specifically samples the mRNA as a complex with eIF4G, rather than simply directing the complex to the cap.

**Supplementary Movie 7. eIF4A recycles eIF4F from non-productive binding locations.** In these ejection experiments, Cy5-eIF4G:E begins the movie bound to the capped (left) and uncapped (right) mRNA constructs. Between frames 2 and 3, the buffer is exchanged removing all free, unbound Cy5-eIF4G:E. However, the buffer it is exchanged into contains eIF4A, eIF4B and ATP. Consequently, on the uncapped mRNA (right), the red spots indicating Cy5-eIF4G:E are rapidly cleared within ~9 seconds from the injection, whereas the Cy5-eIF4G:E remains stably bound at the cap for around a minute (left).
